# Sphingosine 1-phosphate receptor 1 inhibition induces a pro-apoptotic signaling cascade in T cells

**DOI:** 10.1101/2023.08.21.554104

**Authors:** Dhaval Dixit, Victoria M. Hallisey, Ethan Y.S. Zhu, Martyna Okuniewska, Ken Cadwell, Jerry E. Chipuk, Jordan E. Axelrad, Susan R. Schwab

## Abstract

Effective immunity requires a large, diverse naïve T cell repertoire circulating among lymphoid organs in search of antigen. Sphingosine 1-phosphate (S1P) and its receptor S1PR1 contribute by both directing T cell migration and supporting T cell survival. Here, we address how S1P enables T cell survival, and the implications for patients treated with S1PR1 antagonists. Contrary to expectations, we found that S1PR1 limits apoptosis by maintaining the appropriate balance of BCL2 family members via restraint of JNK activity. Interestingly, the same residues of S1PR1 that enable receptor internalization are required to prevent this pro-apoptotic cascade. Findings in mice were recapitulated in ulcerative colitis patients treated with the S1PR1 antagonist ozanimod, and the loss of naïve T cells limited B cell responses. Our findings highlight an unexpected effect of S1PR1 antagonists on the ability to mount immune responses within lymph nodes, beyond their effect on lymph node egress, and suggest both limitations and novel uses of this important class of drugs.

## Introduction

Regulated cell death is an integral process in mammalian development and homeostasis. T cells are often described as “primed” for apoptosis; developing T cells with non-functional or self-reactive T cell receptors must be eliminated(1, 2), and the bulk of effector T cells in an immune response must die to prevent excessive inflammation(3). However, naïve T cells must be long-lived to maintain a diverse repertoire to defend against diverse pathogens, and memory T cells must be long-lived to confer lasting protection(4, 5). A T cell makes critical decisions about whether to live or die at each stage of its life, and the factors that the cell weighs remain incompletely understood.

The signaling lipid sphingosine 1-phosphate (S1P) has been intensively studied in the context of T cell migration(6). The high concentration of S1P in blood and lymph guides T cells out of the low-S1P environment of lymphoid organs, and T cells follow this gradient primarily using S1P receptor 1 (S1PR1). Without circulatory S1P or without S1PR1 on T cells, T cells fail to exit the thymus into blood, and fail to exit lymph nodes (LN) into lymph. We recently found that S1P, acting via S1PR1, also plays an essential role in naïve T cell survival, which interestingly is independent of its role in cell migration (7).

Four drugs that target S1PR1 have been FDA-approved to treat multiple sclerosis, and one has been approved to treat ulcerative colitis. These drugs work in part by blocking pathogenic T cells from exiting LN, thus preventing them from accessing the brain or colon(8, 9). However, some effects of these drugs have been unexplained. Patients treated with drugs targeting S1PR1 respond poorly to SARS-CoV-2 vaccines(10–12), although B cell responses should be initiated within the LN. Moreover, antibody titers have been reported to correlate negatively with time on drug, while the drugs’ effects on cell migration occur within hours of the first dose and rapidly reach a new equilibrium(10–12). Finally, the few studies measuring blood lymphocyte counts in patients who have stopped treatment suggest that some patients do not fully recover cell numbers(13, 14).

We know very little about how S1P supports T cell survival, or about how the effects of genetic blockade of S1P signaling in mice translate to the many patients treated with drugs targeting S1PR1. Here we address these questions. We find that S1PR1 activation restricts JNK phosphorylation, thereby maintaining the appropriate balance of BCL2 family members within the T cell, and in turn limiting apoptosis. Interestingly, the same residues of the S1PR1 C-terminus that enable receptor internalization are required to prevent the apoptotic cascade. Our findings using genetic models were recapitulated in mice treated with the drug fingolimod (FTY720), which targets S1PR1, and in ulcerative colitis patients treated with the drug ozanimod, also an S1PR1 modulator. Using a mouse model, we find that poor antibody responses after prolonged treatment with S1PR1 antagonists may in part reflect loss of the naïve T cell repertoire, pointing to an important role of S1P signaling in regulating T cell responses within the lymph node as well as in non-lymphoid tissues. Our findings simultaneously suggest reasons for caution in use of S1PR1 antagonists and new uses for these drugs.

## Results

### The S1P transporter SPNS2 and S1P receptor 1 prevent naïve T cell death

We set the stage for our work by further testing the role of S1P in promoting naïve T cell survival. We had previously shown that in mice lacking the S1P transporter SPNS2 in lymphatic endothelial cells (*Spns2*^f/f^ *Lyve1-*Cre, *Spns2*Δ^Lyve1^), lymph S1P was lost(7). As expected, naïve T cells were retained in lymph nodes, and were no longer exposed to circulatory S1P(7). Surprisingly, they died at an increased rate compared to T cells in lymph nodes of littermate controls(7). To assess potential developmental defects contributing to T cell death, we asked how inducible deletion of *Spns2* in adults would affect naïve T cell survival. We bred *Spns2*^f/f^ *UBC*-CreERT2 mice, with widespread expression of tamoxifen-inducible Cre(15), and treated adults with tamoxifen 3 weeks before analysis (tamoxifen-treated *Spns2*^f/f^ *UBC*-CreERT2 mice, *Spns2*Δ^UBC^). We measured cell death by flow cytometry, assessing propidium iodide uptake as well as dual staining with a probe for active caspases and with Annexin V (Figure 1A). We found an increased percent of dying cells among naïve T cells in *Spns2*Δ^UBC^ LN compared to tamoxifen-treated littermate controls, as well as a loss of naïve T cells (Figures 1B and S1A-S1B). Consistent with the importance of S1P secretion by SPNS2, we similarly found an increased percent of dying cells among naïve T cells in LN of mice with lymphatic endothelial cell-specific deletion of the two sphingosine kinases that synthesize S1P (*Sphk1^f/f^;Sphk2^-/-^ Lyve1-*Cre, *Sphk*Δ^Lyve1^)(16) (Figure S1C). Finally, we confirmed our previous findings that S1PR1 was required in a cell-intrinsic manner to limit naïve T cell death, using mixed bone marrow chimeras and transfer experiments (below and Ref (7)). These data suggest a model in which S1P secreted via SPNS2 and sensed by T cells via S1PR1 promotes naïve T survival.

**Figure 1:**
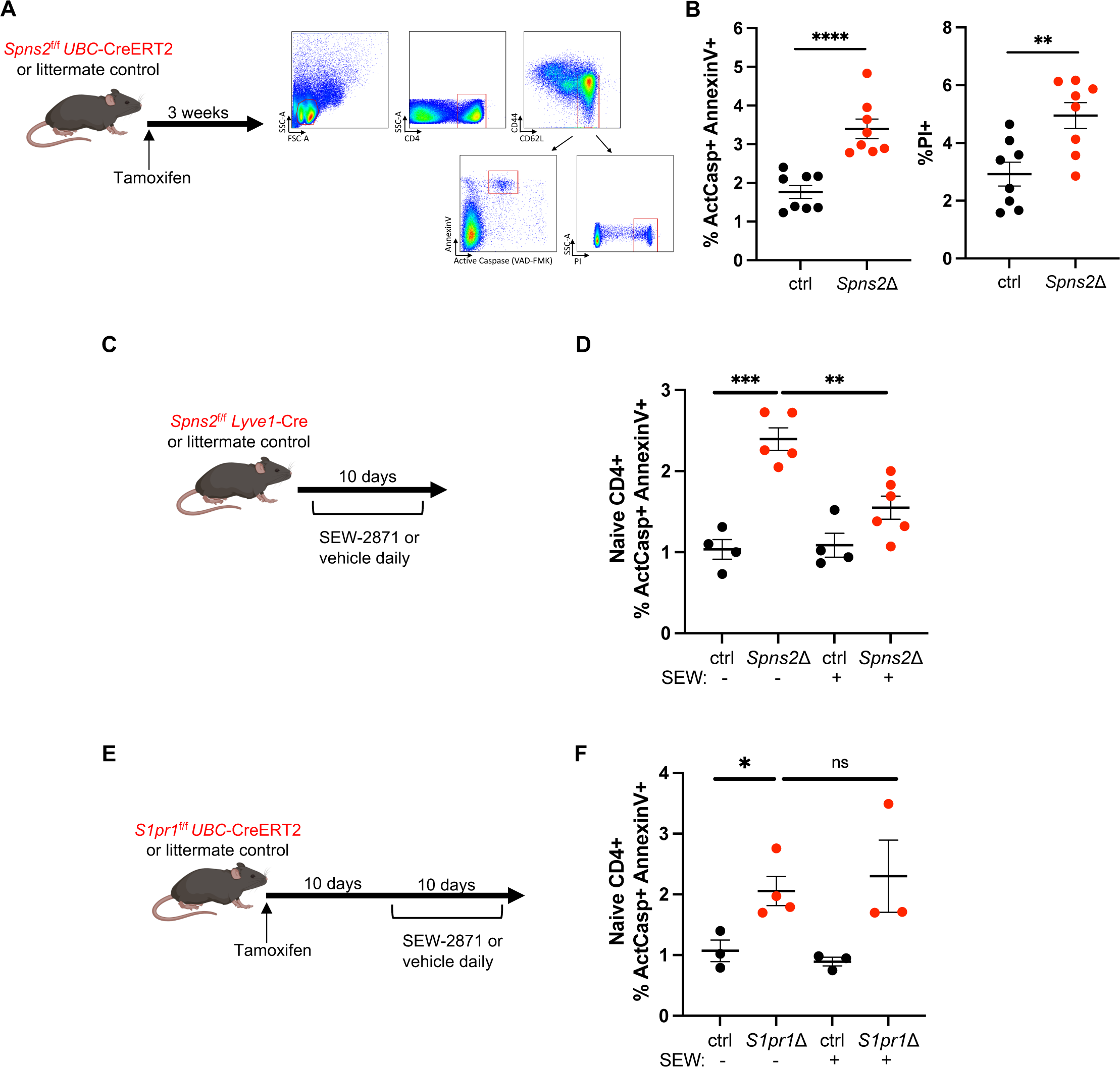
SPNS2-derived S1P and S1PR1 prevent apoptotic death of naïve T cells. A-B) *Spns2*^f/f^ *UBC*-CreERT2 mice and littermate controls were treated with tamoxifen, and 3-4 weeks later LN T cells were analyzed by flow cytometry. (A) Experiment design and gating strategy. (B) Frequency of active caspase (ActCasp)+ AnnexinV+ and propidium iodide (PI)+ cells among naïve CD4+ T cells. Compilation of 4 experiments with n=8 mice for both groups. C-D) *Spns2*^f/f^ *Lyve1*-Cre and littermate controls were treated with 10mg/kg SEW-2871 or vehicle daily. After ten days of treatment, LN T cells were analyzed by flow cytometry. (C) Experiment design. (D) Frequency of ActCasp+ AnnexinV+ cells among naïve CD4+ T cells in LN of *Spns2*Δ mice and littermate controls, with and without SEW-2871 treatment. Compilation of 4 experiments with n=4-6 mice per group. E-F) *S1pr1*^f/f^ *UBC*-CreERT2 mice and littermate controls were treated with 10mg/kg SEW-2871 or vehicle daily for 12 days and treated with tamoxifen on d1 and d2. On d12, LN cells were analyzed by flow cytometry. Frequency of ActCasp+ AnnexinV+ cells among naïve CD4+ T cells in LN of *S1pr1*Δ mice and littermate controls, with and without SEW-2871 treatment. Compilation of 3 experiments with n=3-4 mice per group. Statistical analysis was performed using Student’s t test. *p ≤ 0.05, **p ≤ 0.01, ***p ≤ 0.001, ****p ≤ 0.0001, N.S. non-significant.

### S1P signaling promotes naïve T cell survival independent of exit from LN or uptake of S1P

S1P could promote naïve T cell survival by at least three possible mechanisms: enabling T cell trafficking among or within lymphoid organs, acting as a metabolite or protein co-factor after uptake into the T cell, or activating a signaling pathway downstream of S1PR1 within the T cell.

Our previous work had indicated that S1PR1 did not promote survival by enabling exit from lymph nodes or access to IL7 or peptide-MHC within lymph nodes (Ref. (7) and below). We next considered the hypothesis that S1P uptake was important for T cell survival. S1PR1 is a G-protein-coupled receptor (GPCR), and, like many GPCR, S1PR1 is internalized after binding its ligand(17, 18). Intracellular S1P has been implicated in many processes important for cell survival, from acting as a metabolic intermediate to promoting mitochondrial homeostasis to regulating glycolysis(19–22). To test the role of S1P uptake by S1PR1 in T cell survival, we took advantage of a chemical agonist of S1PR1, SEW-2871, which is structurally distinct from S1P(23). We treated *Spns2*Δ^Lyve1^ mice and littermate controls with SEW-2871 or vehicle daily for 10 days (Figure 1C). Naïve T cells in both *Spns2*Δ^Lyve1^ mice and controls expressed lower levels of surface S1PR1 than naïve T cells in vehicle-treated mice (Figure S1D), consistent with previous findings that SEW-2871 bound the receptor and induced its internalization. As expected, in *Spns2*Δ^Lyve1^ mice, the increased S1PR1 signal in the LN did not restore egress; as expected, in control mice, increased agonism of S1PR1 in the LN disrupted the ligand gradient that would normally guide exit from LN (Figure S1E). We found that SEW-2871-treated *Spns2*Δ^Lyve1^ mice had a reduced frequency of dying naïve T cells compared to vehicle-treated *Spns2*Δ^Lyve^ mice (Figures 1D and S1F). This result suggested that signaling through S1PR1, rather than S1P uptake (or lymph node exit), was important for naïve T cell survival. Importantly, SEW-2871 treatment of *S1pr1*Δ^UBC^ mice did not rescue naïve T cell apoptosis (Figures 1E-1F), indicating that SEW-2871 was working through S1PR1 to promote survival.

### Naïve T cell apoptosis is not associated with changes in canonical signaling pathways downstream of S1PR1

We then considered the possibility that S1PR1 activates a pro-survival signaling pathway within naïve T cells. S1PR1 couples to Gα_i_ and its associated Gβγ subunits(24). Experiments using cultured cells have demonstrated that S1PR1, like many GPCR, can activate AKT and ERK, kinases that promote survival in many contexts(25, 26). Furthermore, Gα_i_ inhibition of adenylate cyclase has been demonstrated in many cell types to prevent cAMP accumulation and hence limit protein kinase A (PKA) activation. Yet little is known about S1PR1-dependent signaling pathways in naïve T cells *in vivo.* In the lymph node, T cells receive inputs from many other sources including cytokine receptors, chemokine receptors, the T cell receptor, and metabolic sensors, and these have also been shown to regulate AKT, ERK, and/or PKA signaling. We therefore asked whether S1PR1’s contribution was limiting in the context of the many other inputs into these pathways in a naïve T cell *in vivo*.

To study S1PR1’s contribution to AKT and/or ERK activation, we measured AKT and ERK phosphorylation in naïve T cells from *Spns2*Δ^Lyve1^ and *S1pr1*Δ^UBC^ mice and littermate controls. We predicted that expression of pAKT or pERK would be decreased in naïve T cells from *Spns2*Δ^Lyve1^ and *S1pr1*Δ^UBC^ mice if S1P signaling were a key input to the pathway. We fixed LN cells rapidly after euthanasia to prevent changes in phosphorylation during cell preparation, and analyzed the cells by flow cytometry. We detected no difference in pAKT or pERK in naïve T cells between *Spns2*Δ^Lyve1^ mice and littermate controls. This could have simply reflected a limitation of the assay, but, surprisingly, naïve T cells from *S1pr1*Δ^UBC^ mice had reproducibly ***increased*** levels of pAKT and pERK compared to littermate controls (Figures 2A-2C). Since activity of these pathways was not decreased in both strains, we concluded that they were unlikely to account for S1P-dependent naïve T cell survival. We were surprised by the divergence between *Spns2*Δ and *S1pr1*Δ T cells in pAKT and pERK levels, and took a detour to investigate the cause; we found that differential expression of the C-type lectin CD69 explained the difference (Supplemental Figure 2).

**Figure 2:**
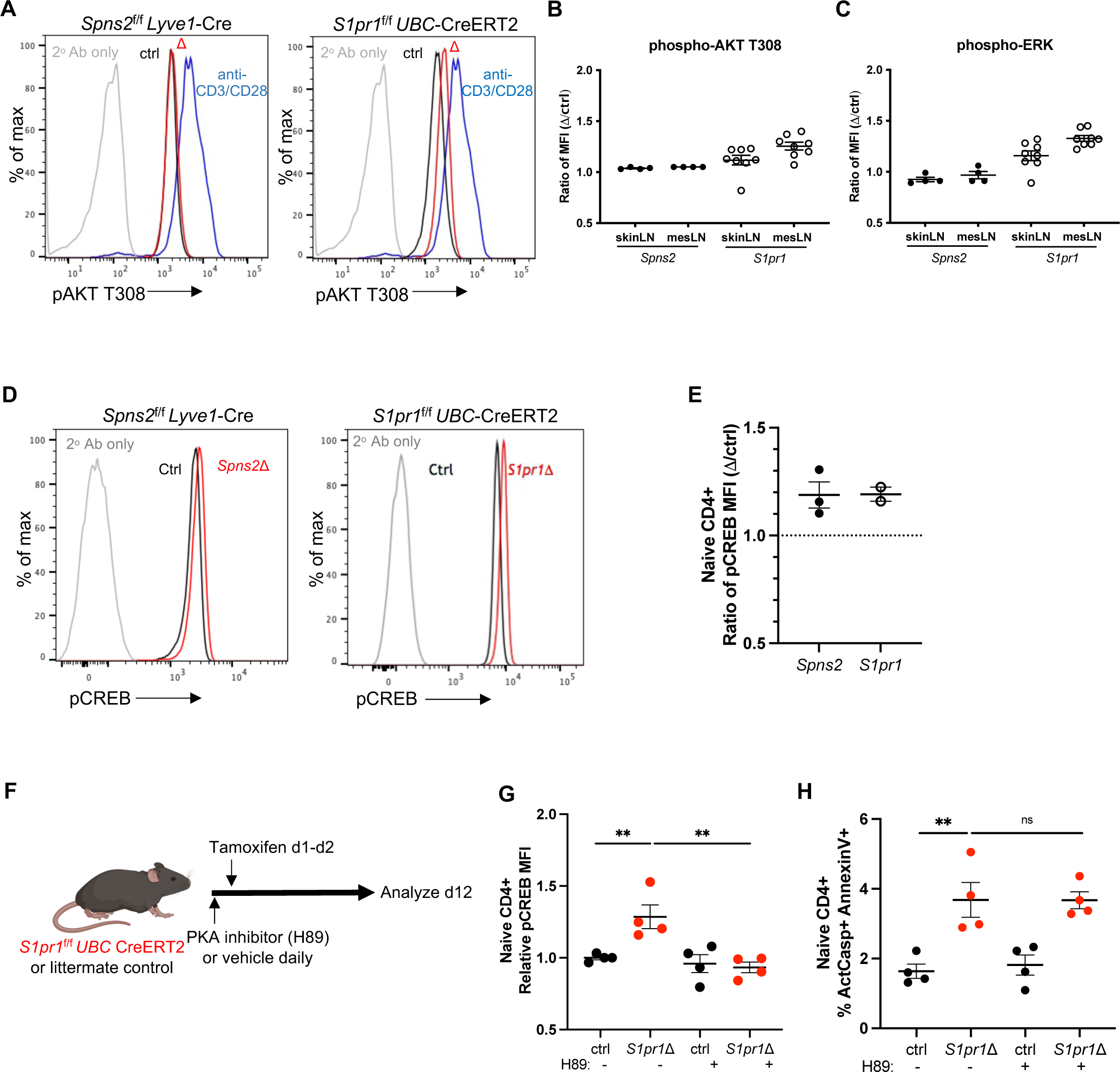
Canonical S1PR1 signaling pathways do not regulate naïve T cell survival. A-B) Phosphorylated AKT T308 (pAKT T308) in T cells from skin-draining LN or mesenteric LN was measured by flow cytometry. (A) Representative histogram of pAKT T308 in naïve CD4+ T cells from the skin-draining LN of (left) a *Spns2*^f/f^ *Lyve1*-Cre mouse and its littermate control, or (right) a *S1pr1*^f/f^ *UBC*-CreERT2 mouse and its littermate control 3-4 weeks after tamoxifen treatment. pAKT T308 in WT CD4 T cells activated *in vitro* served as a comparison (the same comparison was used in both plots). (B) Compilation. Each point represents the ratio of the pAKT T308 geometric mean fluorescence intensity (MFI) in naïve CD4 T cells from a *Spns2*Δ mouse to the pAKT T308 MFI in naïve CD4 T cells from its littermate control, or the ratio of the pAKT T308 MFI in naïve CD4 T cells from a *S1pr1*Δ mouse to the pAKT T308 MFI in naïve CD4 T cells from its littermate control. Compilation of 4 experiments with 4 pairs of mice for *Spns2* and 8 experiments with 8 pairs of mice for *S1pr1*. (C) As in (B), for pERK. D-E) Phosphorylated CREB (pCREB) was measured by flow cytometry. (D) Representative histogram of pCREB in naïve CD4+ T cells from LN of (left) a *Spns2*^f/f^ *Lyve1*-Cre mouse and its littermate control, or (right) a *S1pr1*^f/f^ *UBC*-CreERT2 mouse and its littermate control 3-4 weeks after tamoxifen treatment. (E) Compilation of pCREB expression. Each point represents the ratio of pCREB geometric MFI between a *Spns2*Δ or *S1pr1*Δ mouse and its littermate control. Compilation of 2-3 experiments with n=2-3 per group. F-H) *S1pr1*^f/f^ *UBC*-CreERT2 mice and littermate controls were treated with 20mg/kg H-89 (PKA inhibitor) or vehicle daily starting on d0. They were treated with tamoxifen to delete *S1pr1* on d1 and d2. LN T cells were analyzed by flow cytometry on d12. (F) Experiment design. (G) Relative pCREB expression in naïve CD4+ T cells. Each point represents the geometric MFI value in one individual mouse divided by the mean geometric MFI value in control mice treated with vehicle in that experiment. (H) Frequency of ActCasp+ AnnexinV+ cells among naïve CD4+ T cells in LN of the indicated mice. Compilation of 3 experiments, with 4 mice in each group. Statistical analysis was performed using Student’s t test. *p ≤ 0.05, **p ≤ 0.01, ***p ≤ 0.001, ****p ≤ 0.0001, N.S. non-significant.

We also tested whether Gα_i_ inhibition of cAMP-PKA was important for S1P-dependent naïve T cell survival. Activated PKA can phosphorylate the transcription factor CREB(27). We found that naïve T cells from both *Spns2*Δ and *S1pr1*Δ mice had slightly increased phosphorylated CREB, suggesting that S1PR1’s input was limiting for inhibition of the cAMP-PKA-CREB pathway (Figures 2D-2E). To test whether this pathway was important in naïve T cell survival, we treated *S1pr1*Δ mice daily with a PKA inhibitor, H89(28) (Figure 2F). Although the inhibitor reduced CREB phosphorylation, it did not affect the frequency of dying naïve T cells, suggesting that a PKA-dependent pathway did not regulate S1P-dependent naïve T cell survival (Figures 2G-2H).

Overall, these data suggest that canonical signaling pathways downstream of S1PR1 do not regulate S1P-dependent naïve T cell survival.

### S1P signaling promotes survival by regulating the balance of BCL2 family proteins

We next investigated whether loss of S1PR1 signaling resulted in changes in proximal regulators of cell death; we hoped to use this information to work our way back to the receptor.

Analysis of RNA and protein indicated that the balance of BCL2 family members was altered upon loss of S1P signaling. Using RNA-Seq, we compared transcripts of *S1pr1*Δ^UBC^ and littermate control naïve T cells, isolated from mixed bone marrow chimeras in which *S1pr1*^f/f^*UBC*-CreERT2 and control bone marrow was used to reconstitute WT hosts. We also compared transcripts of naïve T cells from chimeras in which WT bone marrow was used to reconstitute either *Spns2*Δ^Lyve1^ or littermate control hosts. Two genes within the BCL2 family were differentially expressed upon loss of S1P signaling: the anti-apoptotic gene *Bcl2* (BCL2) was down-regulated (more consistently in SPNS2-deficient than S1PR1-deficient T cells), and the pro-apoptotic gene *Bbc3* (PUMA) was up-regulated (Figure S3A). Using flow cytometry to measure intracellular protein, we found that naïve T cells from *Spns2*Δ and *S1pr1*Δ mice had an approximately 20-25% reduction in BCL2 staining compared to littermate controls (Figures 3A-3B and S3C). Furthermore, we detected an approximately 20-25% increase in PUMA staining (Figures 3C-3D and S3D). Ultimately, apoptosis is executed by effectors, either BAX or BAK. Naive T cells from *Spns2*Δ and *S1pr1*Δ mice had an approximately 30% increase in BAX staining compared to controls (Figures 3E-3F and S3E). Because T cells are thought to be “primed” for apoptosis, we thought it plausible that this imbalance of BCL2 family members might trigger cell death via the mitochondrial pathway of apoptosis.

**Figure 3:**
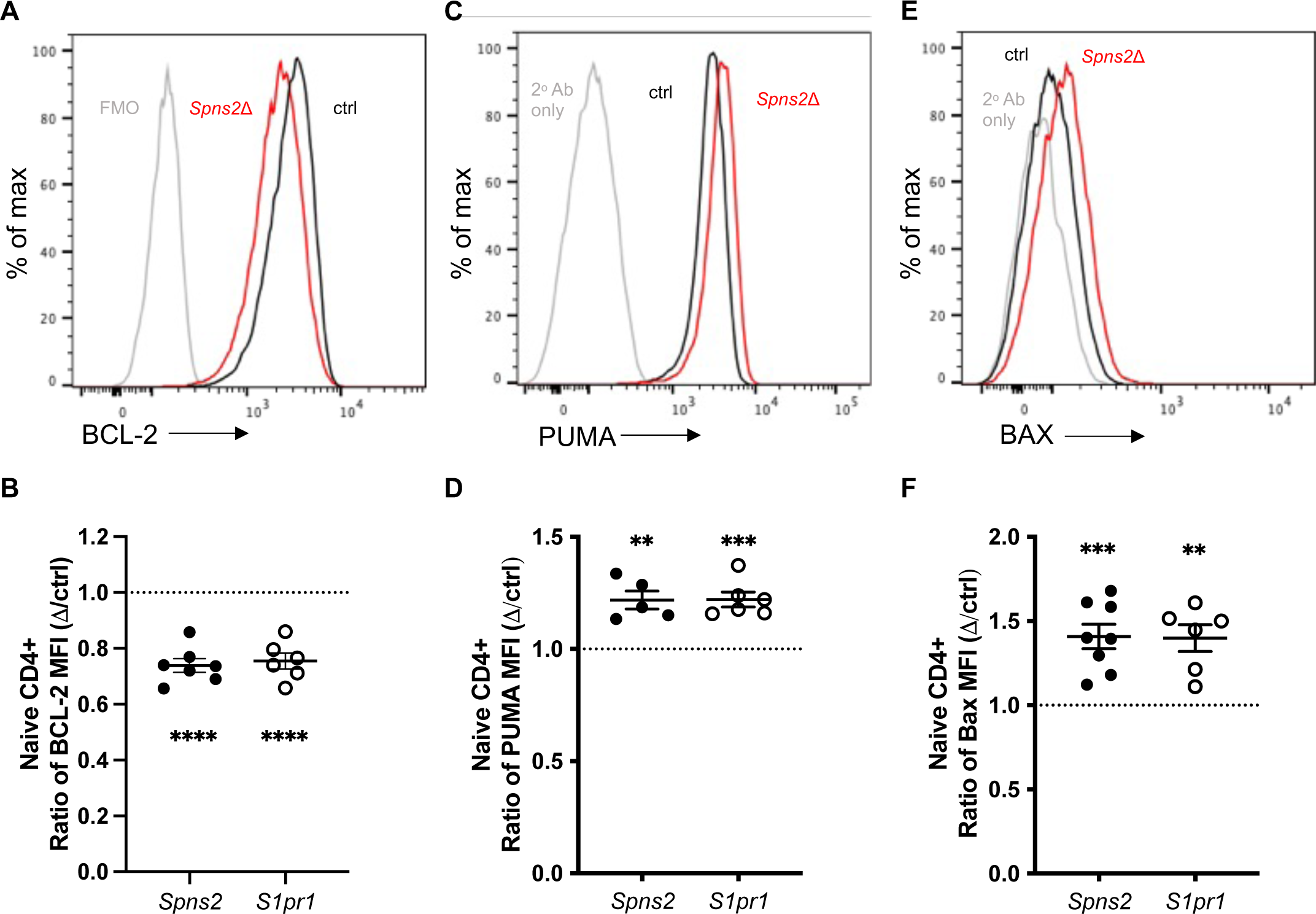
BCL-2 family proteins regulate S1PR1-dependent naïve T cell survival. Expression of BCL-2 family members by naïve CD4+ T cells from LN of *Spns2*Δ or *S1pr1*Δ mice and littermate controls, analyzed by flow cytometry. (A-B) Representative histogram from a *Spns2*^f/f^ *Lyve1*-Cre mouse and its littermate control, and compilation of BCL-2 expression. Each point represents the ratio of BCL-2 geometric MFI between a *Spns2*Δ or *S1pr1*Δ mouse and its littermate control (or the mean of littermate controls if there were more than one). Compilation of 5-6 experiments with n=5-7 per group. (C-D) As in (A-B) for PUMA. Compilation of 5-6 experiments with n=5-6 pairs per group. (E-F) As in (A-B) for BAX. Compilation of 6 experiments with n=5-8 pairs per group. Statistical analysis was performed using Student’s t test, comparing (KO MFI)/(mean ctrl MFI) to (ctrl MFI)/(mean ctrl MFI). *p ≤ 0.05, **p ≤ 0.01, ***p ≤ 0.001, ****p ≤ 0.0001, N.S. non-significant.

To assess whether this imbalance of BCL2 family members resulted in death upon loss of S1P signaling, we tested three predictions. First, we predicted that if *S1pr1*Δ T cells had a physiologically relevant imbalance of BCL2 family members, providing an additional stress that activated the mitochondrial pathway of apoptosis would lead to increased death of *S1pr1*Δ T cells compared to controls. We co-transferred *S1pr1*Δ and littermate control naïve T cells at a 1:1 ratio into WT recipients. One day after transfer, we treated the recipients with the BCL2-specific inhibitor ABT-199(29), and one day after ABT-199 treatment, we enumerated transferred cells in LN (Figure 4A). We found that ABT-199 treatment led to a significant reduction in the ratio of *S1pr1*Δ to control T cells (Figure 4B). Similar *in vivo* experiments using dexamethasone or irradiation also led to a reduction in the ratio of *S1pr1*Δ to control T cells, suggesting that *S1pr1*Δ T cells are more susceptible than controls to pro-apoptotic stresses (Figures S3F-S3G). ABT-199 treatment *ex vivo* also led to increased rates of cell death among *S1pr1*Δ cells compared to controls (Figures 4C-4D). By contrast, *ex vivo* treatment with Fas ligand led to similar rates of cell death among *S1pr1*Δ T cells and controls (Figures 4C and 4E), suggesting that *S1pr1*Δ T cells were not more susceptible to death via the extrinsic pathway of apoptosis.

**Figure 4:**
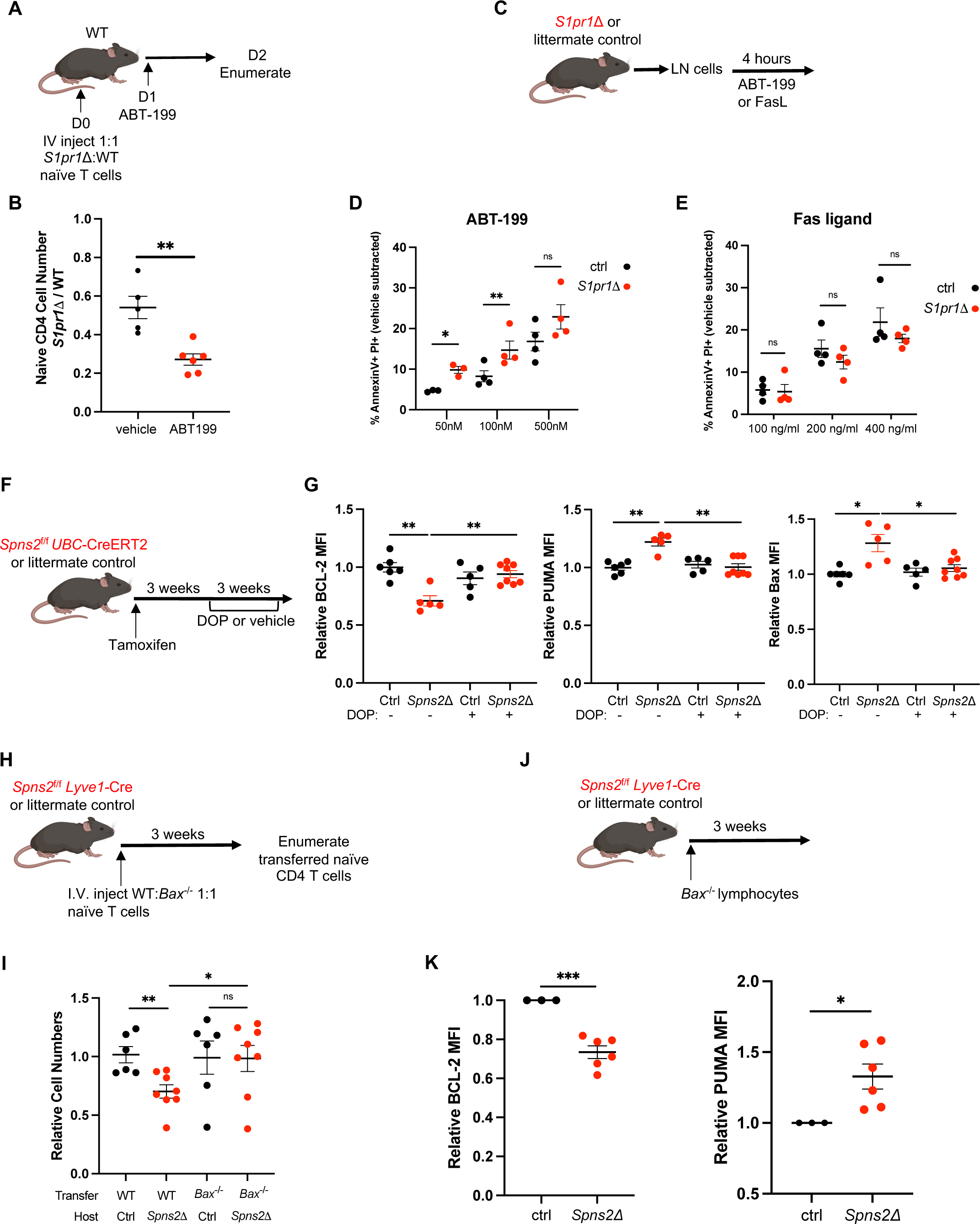
BCL-2 family proteins regulate S1PR1-dependent naïve T cell survival. A-B) *S1pr1*Δ and littermate control lymphocytes were co-transferred (1:1 by naïve CD4 T cell counts) intravenously into WT recipients. The transferred cells were labeled with CellTrace Violet or CellTrace Yellow, and dyes were swapped between experiments. 24 hours later, recipients were treated with ABT-199. 24 hours after ABT-199 treatment, dye-labeled naïve CD4 T cells were enumerated in LN. (A) Experiment model. (B) Ratio of the number of naïve CD4+ T cells from *S1pr1*Δ vs its littermate control recovered in recipients, with ABT-199 or control treatment. Compilation of 3 experiments with n=5-6 mice per group. C-E) Lymphocytes from LN of *S1pr1*Δ or littermate control mice were treated with the indicated concentrations of (D) ABT-199 or (E) Fas ligand *ex vivo* for 4 hours. The frequency of AnnexinV+ PI+ cells among naïve CD4+ T cells was measured by flow cytometry. Each point represents the mean of two technical replicates, minus the mean frequency of AnnexinV+ PI+ cells in two vehicle-treated technical replicates. Compilation of 3 experiments with n=4 mice per group. F-G) *Spns2*^f/f^ *UBC*-CreERT2 mice and littermate controls were treated with tamoxifen. At least 3 weeks later, mice were treated with 30mg/l DOP and 10mg/l sucrose, or sucrose alone, in the drinking water. After 3 weeks of DOP treatment, LN T cells were analyzed by flow cytometry. (F) Experiment strategy. (G) Relative BCL-2, PUMA, and BAX expression in naïve CD4+ T cells. Each point represents the geometric MFI value in one individual mouse divided by the mean MFI value in control mice treated with vehicle in that experiment. Compilation of 4 experiments with n=5-8 mice per group. H-I) *Bax*^-/-^ and WT littermate control lymphocytes were co-transferred (1:1 by naïve CD4 T cell counts) intravenously into *Spns2*Δ mice and littermate controls. 21 days later, naïve CD4 T cells from skin-draining and mesenteric LN were enumerated. The transferred cells were labeled with CellTrace Violet or CellTrace Yellow, and dyes were swapped between experiments. (H) Experiment design. (I) Relative numbers of transferred naïve CD4+ T cells. Each point represents the count in one individual mouse divided by the mean number of WT cells in the control LN in that experiment. Compilation of 3 experiments with n=6-8 mice per group. J-K) *Bax*^-/-^ lymphocytes were transferred intravenously into *Spns2^f^*^/f^ *Lyve1*-Cre mice or littermate controls. The transferred cells were labeled with CellTrace Violet. 21 days later, dye-labeled LN naïve CD4+ T cells were analyzed by flow cytometry. (J) Experiment strategy. (K) Compilation of BCL-2 and PUMA expression by *Bax*^-/-^ naïve T cells. Each point represents the geometric MFI value in one individual mouse divided by the mean geometric MFI value of control mice in that experiment. Compilation of 3 experiments with n=3-6 mice per group. Statistical analysis was performed using Student’s t test. *p ≤ 0.05, **p ≤ 0.01, ***p ≤ 0.001, ****p ≤ 0.0001, N.S. non-significant.

Second, we predicted that if loss of S1P signaling led to an imbalance of BCL2 family members, then increasing T cell exposure to S1P in *Spns2*Δ mice should restore the balance. To test this, we took advantage of the finding that the enzyme S1P lyase maintains low levels of S1P in LN(18). We treated *Spns2*Δ^UBC^ and control mice with the S1P lyase inhibitor 4’deoxyhydropyridoxine (DOP) for 3 weeks (Figure S4A). As expected from previous studies(*12*), DOP treatment of *Spns2*Δ or control mice increased T cell exposure to S1P within LN, indicated by internalization of S1PR1 (Figure S4B). As expected, in *Spns2*Δ^UBC^ mice, the increased LN S1P did not restore exit from LN; in control mice, the increased LN S1P disrupted the gradient that would normally guide cells out of LN and limited exit (Figure S4C). As expected, DOP-treated *Spns2*Δ^UBC^ mice had a reduced frequency of dying naïve T cells compared to vehicle-treated *Spns2*Δ^UBC^ mice (Figures S4D-S4E). We assessed how DOP affected the levels of BCL2 family members in naïve T cells. We found that DOP treatment restored BCL2, PUMA, and BAX levels in naïve T cells from *Spns2*Δ mice to similar levels as controls, consistent with the hypothesis that S1P signaling maintains the balance of BCL2 family members in naïve T cells (Figures 4F-4G).

Last, we predicted that the imbalance of BCL2 family members would be the upstream cause of apoptosis in mice lacking S1P signaling. To test this, we took advantage of *Bax*^-/-^ T cells(30), which lack one of the two pore-forming proteins that execute the mitochondrial pathway of apoptosis. We co-transferred T cells from *Bax*^-/-^ mice and WT littermates into *Spns2*Δ^Lyve1^ mice and littermate controls. 3 weeks later, we enumerated the transferred cells (Figure 4H). Transferred WT T cells decreased about 30% in number in LN of *Spns2*Δ^Lyve1^ mice compared to WT T cells in LN of controls. By contrast, *Bax*^-/-^ T cells in LN of *Spns2*Δ^Lyve1^ mice were found in similar numbers to *Bax*^-/-^ or WT T cells in LN of controls (Figure 4I). Although *Bax*^-/-^ T cells survived better in Spns2Δ^Lyve1^ mice than control mice, *Bax*^-/-^ T cells transferred into *Spns2*Δ mice still had lower levels of BCL2 and higher levels of PUMA than *Bax*^-/-^ T cells transferred into controls (Figures 4J-4K and S4F). This result suggested that the imbalance of these proteins preceded death.

These data suggest that lack of S1P signaling leads to an imbalance in BCL2 family proteins, which promotes naïve T cell apoptosis. Of course, additional changes in BCL2 family members beyond those that we measured may also contribute to this imbalance.

### S1P signaling regulates the balance of BCL2 family members by restraining JNK activation

The expression and function of BCL2 family members are regulated by multiple pathways, including JNK signaling(31). Prolonged JNK activation can promote apoptosis(32). We therefore tested whether JNK was activated in cells lacking S1P signaling. We measured phosphorylation of JNK and one of its downstream targets, the transcription factor cJun, by flow cytometry. Naïve T cells from LN of both *Spns2*Δ and *S1pr1*Δ mice had reproducibly increased expression of phosphorylated JNK1/2 (pJNK) and phosphorylated cJun (pcJun) compared to controls (Figures 5A-5B, S5A-S5B). JNK can be activated by the kinases MKK4 and MKK7(32), and we detected increased phosphorylated MKK7 (pMKK7) in naïve T cells from *Spns2*Δ and *S1pr1*Δ mice (Figures 5C and S5C).

**Figure 5:**
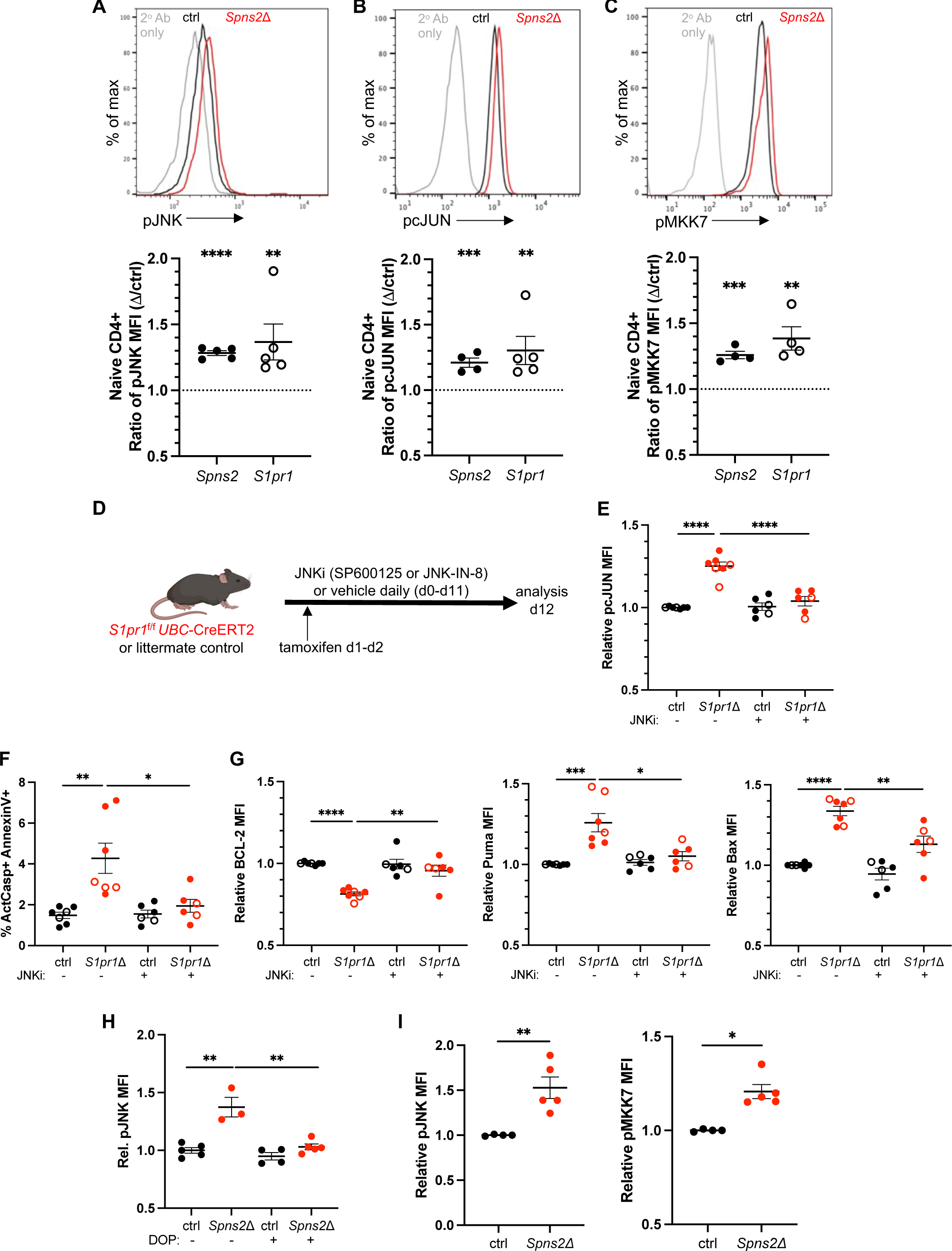
JNK signaling regulates S1P-dependent naïve T cell survival. A-C) Expression of phosphorylated JNK pathway proteins in naïve CD4+ T cells from LN of *Spns2*Δ or *S1pr1*Δ mice and littermate controls, analyzed by flow cytometry. (A) Representative histogram from a *Spns2*^f/f^ *Lyve1*-Cre mouse and its littermate control and compilation of pJNK expression. Each point represents the ratio of pJNK geometric MFI between a *Spns2*Δ or *S1pr1*Δ mouse and its littermate control (or the mean of littermate controls, if more than one were available). Compilation of 4-5 experiments with n=5-6 mice per group. (B) As in (A) for pcJUN. Compilation of 4-5 experiments with n=5-6 pairs per group. (C) As in (A) for pMKK7. Compilation of 4 experiments with n=4-5 mice per group. D-G) Starting on day 0, *S1pr1^f^*^/f^ *UBC*-CreERT2 and littermate control mice were treated daily with either 15 mg/kg body weight SP-600125 (filled circles) or 20 mg/kg body weight JNK-IN-8 (open circles) or vehicle intraperitoneally. On days 1 and 2, the mice were treated with tamoxifen. On day 12, naïve CD4+ T cells in LN were analyzed. Relative values represent expression in one individual mouse divided by the mean value of control mice treated with vehicle in that experiment. (D) Experiment strategy. (E) Relative pcJUN expression. (F) Frequency of ActCasp+ AnnexinV+ among naïve CD4+ T cells. (G) Relative BCL-2, PUMA, and BAX expression. Compilation of 6 experiments (4: SP600125; 2: JNK-IN-8) with n=6-7 mice per group. (H) *Spns2*^f/f^ *UBC*-CreERT2 mice and littermate controls were treated with tamoxifen. 3 weeks later, mice were treated with 30mg/l DOP and 10mg/l sucrose, or sucrose alone, in drinking water. After 3 weeks of DOP treatment, naïve CD4+ T cells in LN were stained for pJNK. Relative values represent expression in one individual mouse divided by the mean value of control mice treated with vehicle in that experiment. Compilation of 3 experiments with n=3-5 mice per group. I) *Bax*^-/-^ lymphocytes were transferred intravenously into *Spns2^f^*^/f^ Lyve1-Cre mice or littermate controls. The transferred cells were labeled with CellTrace Violet. 21 days later, dye-labeled naïve CD4+ T cells in LN were analyzed by flow cytometry. Compilation of relative pJNK and pMKK7 expression. Relative values represent the geometric MFI value in one individual mouse divided by the mean geometric MFI value of control mice in that experiment. Compilation of 2 experiments with n=4-5 mice per group. Statistical analysis was performed using Student’s t test. *p ≤ 0.05, **p ≤ 0.01, ***p ≤ 0.001, ****p ≤ 0.0001, N.S. non-significant.

To assess whether JNK activation resulted in the imbalance of BCL2 family members and increased apoptosis in the absence of S1P signaling, we tested three predictions. First, we predicted that decreasing JNK activity would restore levels of BCL2 family members and T cell survival. We treated *S1pr1*Δ mice with one of two structurally distinct JNK inhibitors, SP600125(33) or JNK-IN-8(34), or vehicle for 10 days (Figure 5D). JNK inhibition effectively reduced levels of pcJun (Fig. 5E). We found that JNK inhibition decreased the frequency of dying naïve T cells in LN (Figure 5F), and that this was accompanied by increased BCL2 and decreased PUMA and BAX (Figure 5G).

Second, we predicted that if loss of S1P signaling led to JNK activation, then increasing T cell exposure to S1P in *Spns2*Δ mice should reduce pJNK. We treated *Spns2*Δ^UBC^ and littermate control mice with DOP for 3 weeks to increase LN S1P. We found that DOP treatment restored pJNK in naïve T cells from *Spns2*Δ mice to levels similar to controls (Figure 5H).

Last, we predicted that JNK activation would be the upstream cause of apoptosis in mice lacking S1P signaling. To test this, we again took advantage of *Bax*^-/-^ T cells, which partly recovered in numbers in *Spns2*Δ hosts. We transferred *Bax*^-/-^ lymphocytes into *Spns2*Δ mice or controls and measured levels of pJNK and pMKK7 3 weeks later. *Bax*^-/-^ T cells in *Spns2*Δ mice had elevated pJNK and pMKK7 compared to *Bax*^-/-^ T cells in controls (Figures 5I and S5D), suggesting that JNK activation preceded death.

These results suggest that loss of S1P signaling results in JNK activation, which in turn alters the balance of BCL2 family members and leads to apoptosis.

### JNK activation is not explained by loss of mitochondria

JNK can be activated by many types of cellular stress(32). We had previously observed that *S1pr1*Δ naïve T cells had decreased mitochondrial content(7), and we asked whether altered mitochondria could be the key stressor. We began by analyzing mitochondria in naïve T cells by electron microscopy (Figure S5E). We found that mitochondria in *S1pr1*Δ naïve T cells had decreased cross-sectional area compared to mitochondria in controls, and that *S1pr1*Δ naïve T cells had a slightly increased number of mitochondria per cell compared to controls (Figures S5F-I). However, we did not detect differences in cross-sectional area of mitochondria or number of mitochondria in naïve T cells from *Spns2*Δ mice compared to controls (Figures S5F-I). We did not observe striking differences in mitochondrial cristae organization between either *S1pr1*Δ mice or *Spns2*Δ mice and their respective controls. While the mitochondrial changes are fascinating, they are unlikely to explain JNK activation because they are divergent between *S1pr1*Δ and *Spns2*Δ T cells.

### JNK activation correlates with loss of S1PR1 internalization

To address how S1P/S1PR1 restrain JNK, we next turned to our RNA-Seq datasets. Interestingly, we found dysregulation of genes related to endocytosis and the actin cytoskeleton in both T cells lacking S1PR1 and T cells from SPNS2-deficient mice (Figure S3B). One striking quality of S1PR1 is that it is extremely susceptible to ligand-induced endocytosis by concentrations of S1P encountered *in vivo*. During a T cell’s continual trafficking among lymphoid organs, S1PR1 is cyclically lost from the cell surface in the high-S1P environment of blood and lymph, and returned to the cell surface in the low-S1P environment of lymphoid organs(35). We therefore considered the possibility that the constant endocytosis of the receptor itself was important for cell physiology, perhaps by affecting the actin cytoskeleton. Our data so far were consistent with a requirement for S1PR1 internalization in T cell survival. In *Spns2*Δ mice, where naïve T cells die, T cells in the LN maintain very high levels of surface S1PR1 due to lack of S1P exposure. In *Spns2*Δ mice treated with DOP or SEW-2871, naïve T cell survival is restored, and surface S1PR1 levels are reduced by ligand-induced internalization.

To test whether internalization of S1PR1 was important for naïve T cell survival, we took advantage of the knowledge that the kinase GRK2 induces S1PR1 internalization by phosphorylating the receptor’s C-terminal tail and enabling β-arrestin to bind the receptor(36, 37). In *Grk2*^f/-^ CD4-Cre mice, naïve T cells maintain aberrantly high levels of S1PR1 in blood and are present at reduced numbers in blood and secondary lymphoid organs(37). We generated bone marrow chimeras by reconstituting WT mice with congenically marked *Grk2*^f/-^ CD4-Cre (*Grk2*Δ) or *Grk2*^f/+^ CD4-Cre (Ctrl) bone marrow (Figure 6A). Eight weeks after reconstitution, we confirmed increased S1PR1 levels on naïve CD4+ T cells in blood (Figures S5J-S5K). We found that LN of *Grk2*Δ mice had an increased frequency of dying naïve T cells, accompanied by increased JNK activation and an imbalance of BCL2 family members, with the exception of similar BCL2 expression between *Grk2*Δ and control naïve CD4 T cells (Figures 6B-6D). These results are largely consistent with the possibility that deficient S1PR1 internalization impairs naïve T cell survival, but interpretation is complicated by pleotropic functions of GRK2.

**Figure 6:**
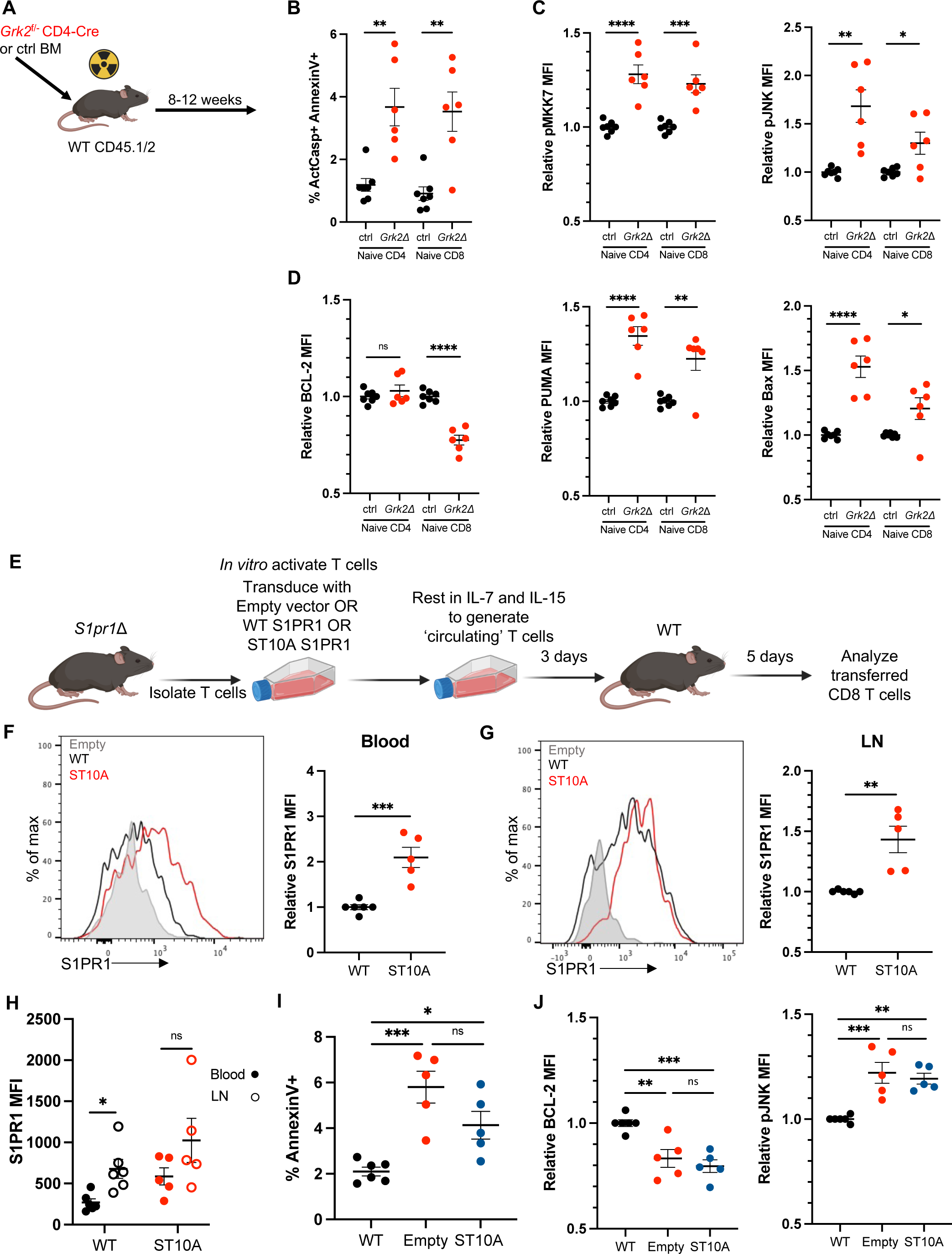
Loss of S1PR1 internalization contributes to apoptosis. A-D) CD45.1/2 WT mice were lethally irradiated and reconstituted with either *Grk2*^f/-^ *Cd4*-Cre (*Grk2*Δ) or *Grk2*^f/+^ *Cd4*-Cre (ctrl) bone marrow. 8-12 weeks after bone marrow transfer, naïve T cells were analyzed. (A) Experiment design. (B) Frequency of ActCasp+ AnnexinV+ cells among naïve CD4+ T cells in LN. (C) Relative MFI values for p-MKK7 and p-JNK and (D) for BCL-2, PUMA, and BAX. Relative values represent expression in one individual mouse divided by the mean value of littermate control mice in that experiment. Compilation of 4 experiments with n=6-7 mice per group. E-J) T cells were isolated from LN of *S1pr1*Δ mice, activated *in vitro* with anti-CD3/CD28, and then retrovirally transduced with vector encoding either IRES-GFP (empty vector), WT S1PR1-IRES-GFP, or ST10A S1PR1-IRES-GFP. Transduced T cells were cultured for three days in IL-7/IL-15 media to generate predominantly CD8+ ‘central memory-like’ T cells. These T cells were transferred into WT recipients and, 5 days later, transferred CD8+ T cells were analyzed. (E) Experiment design. (F-G) Representative histogram and compilation of relative surface S1PR1 expression on cells in (F) blood and (G) LN. Relative surface S1PR1 represents MFI on cells in one individual mouse divided by the mean MFI on WT S1PR1-transduced T cells in that experiment. (H) Compilation of absolute S1PR1 MFI values on the indicated cells. (I) Frequency of AnnexinV+ cells among CD8+ GFP+ T cells in LN. Compilation of 5 experiments with n=5-6 per group. (J) Relative geometric MFI values for BCL-2 and p-JNK in CD8+ GFP+ T cells in LN. Relative values represent expression in one individual mouse divided by the mean value of littermate control mice in that experiment. Compilation of 5 experiments with n=5-6 per group. Statistical analysis was performed using Student’s t test for Figures 6B-D, 6F-H and one-way ANOVA with Tukey’s multiple comparisons test for Figures 6I-J. *p ≤ 0.05, **p ≤ 0.01, ***p ≤ 0.001, ****p ≤ 0.0001, N.S. non-significant.

To study S1PR1 internalization more specifically, we manipulated the C-terminal tail of S1PR1, which has a series of serine and threonine residues that are phosphorylated by GRK2. Replacement of 5 serines in the C-terminal tail with alanine (S351A, S353A, S355A, S358A, S359A)(38) or of 2 serines and 1 threonine in the C-terminal tail with alanine (T371A, S374A, S375A)(37) had been shown to slow S1PR1 internalization, but in both cases surface S1PR1 remained undetectable on T cells in blood and no abnormalities in T cell numbers were reported. To more completely limit internalization, we mutated S1PR1 to substitute alanine for nine serines and one threonine in the C-terminal tail (ST10A). Naïve *S1pr1*Δ T cells were activated in culture; transduced with either empty vector, S1PR1-WT, or S1PR1-ST10A; and then rested for 3 days in IL7 and IL15 to generate CD8+ T cells with central-memory properties(39). Empty vector, S1PR1-WT, or S1PR1-ST10A-expressing *S1pr1*Δ T cells were transferred into recipients. 4-5 days later, T cells from blood and LN were analyzed (Figure 6E). S1PR1-ST10A-transduced *S1pr1*Δ T cells had higher levels of surface S1PR1 compared to S1PR1-WT-transduced *S1pr1*Δ T cells in blood and LN (Figures 6F-6G). S1PR1-ST10A-transduced T cells in LN still had higher surface S1PR1 than S1PR1-ST10A-transduced T cells in blood, although the difference did not reach statistical significance, suggesting that the internalization defect was substantial but not complete (Figure 6H). While S1PR1-WT-transduced *S1pr1*Δ T cells had lower rates of apoptosis than empty vector-transduced *S1pr1*Δ T cells, S1PR1-ST10A-transduced T cells had similarly high rates of apoptosis to empty vector-transduced *S1pr1*Δ T cells (Figure 6I). Similarly, while S1PR1-WT-transduced *S1pr1*Δ T cells had lower levels of pJNK and higher levels of BCL2 than empty vector-transduced *S1pr1*Δ T cells, S1PR1-ST10A-transduced T cells had similar levels of pJNK and BCL2 to empty vector-transduced *S1pr1*Δ T cells (Figure 6J).

Taken together, these data suggest that phosphorylation of the C-terminal tail of S1PR1 regulates both receptor internalization and T cell survival, consistent with the possibility that endocytosis of S1PR1 restrains JNK activation, maintains the balance of BCL2 family members, and prevents apoptosis.

### Chronic FTY720 treatment recapitulates apoptosis

S1PR1 modulators have had remarkable clinical efficacy in treating multiple sclerosis as well as ulcerative colitis. FTY720, the most commonly used drug targeting S1PR1, acts as an agonist of S1PR1 in the short-term but as a functional antagonist of S1PR1 in the long-term in many contexts. After binding S1PR1 and inducing an acute signal through the receptor, FTY720 induces S1PR1 internalization and degradation, mimicking genetic loss of *S1pr1*(40). By contrast, S1P or SEW-2871 induce S1PR1 signaling and internalization, but S1PR1 is then shuttled back to the cell surface where it can signal again(40). Recent studies measuring blood lymphocyte counts in patients who have stopped FTY720 treatment suggest that some patients do not fully recover cell numbers many months after treatment cessation(13, 14). We hypothesized that FTY720 may not only block lymphocyte circulation but also induce naïve T cell death.

To test this, we treated WT mice with FTY720 or vehicle control for 3 weeks (Figure 7A). We found an increased frequency of dying cells among naïve CD4 T cells in LN of FTY720-treated mice compared to controls, accompanied by increased pJNK and pMKK7, decreased BCL2, and increased PUMA and BAX (Figures 7B-7D). Although FTY720 has pleotropic effects, including blockade of thymic exit, these data are consistent with FTY720 acting as a functional antagonist of S1PR1 in the context of T cell death.

**Figure 7:**
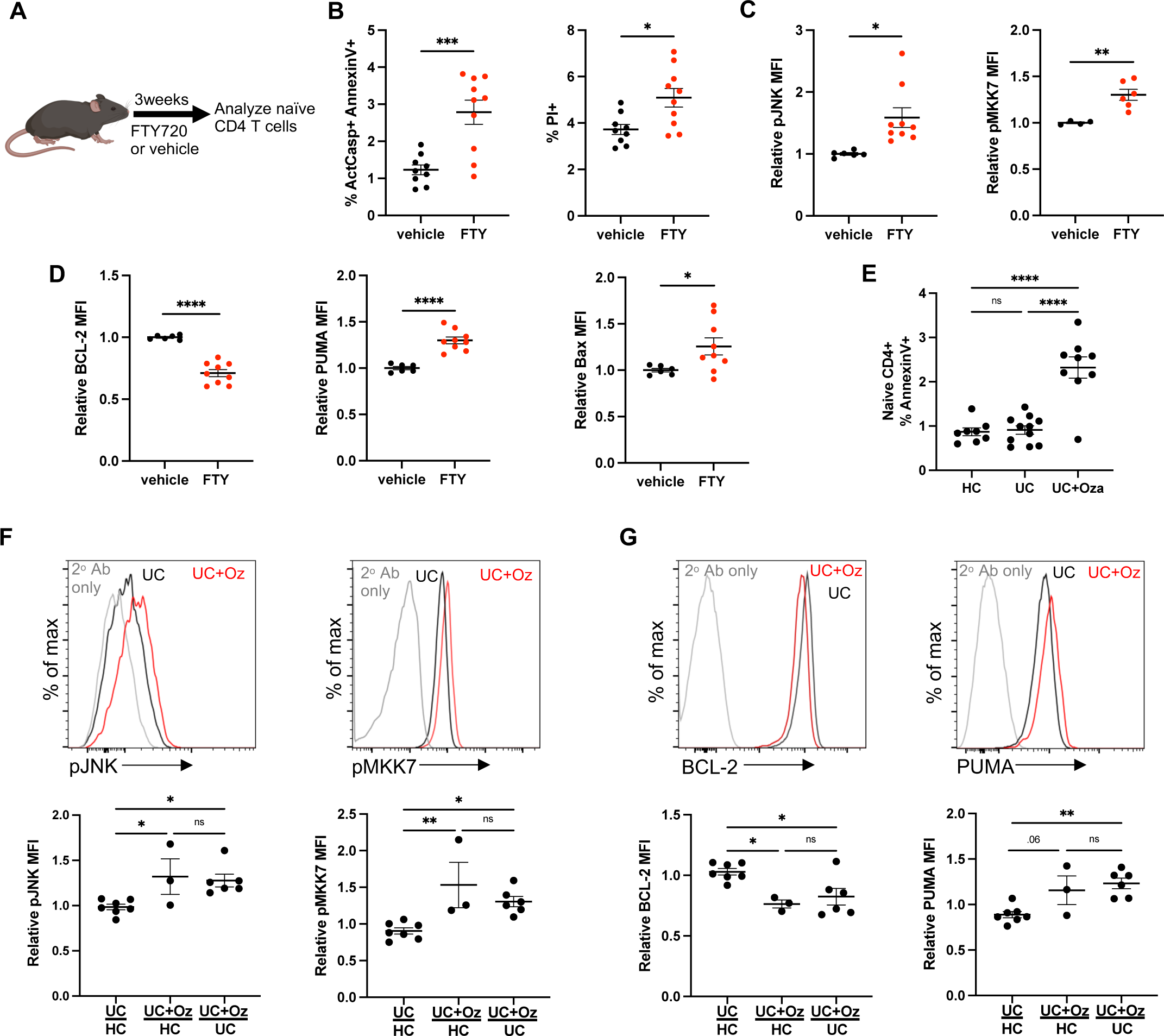
S1PR1 modulators impair T cell survival in mice and humans. A-D) C57BL/6 mice were treated with 1mg/kg FTY720 or vehicle every other day for three weeks, and naïve CD4+ T cells in LN were analyzed by flow cytometry. (A) Experiment design. (B) Frequency of ActCasp+ AnnexinV+ and PI+ cells among naïve CD4+ T cells in LN. Compilation of 3 experiments with n=9-10 mice per group. (C) Relative geometric MFI values for pMKK7 and pJNK. Relative values represent expression in one individual mouse divided by the mean expression in littermate control mice in that experiment. pMKK7: compilation of 2 experiments with n=4-6 per group. pJNK: compilation of 3 experiments with n=6-9 per group. (D) Relative geometric MFI values for BCL-2, PUMA, and BAX. Compilation of 3 experiments with n=6-9 per group. E-G) PBMCs were isolated from blood samples of UC patients treated with ozanimod (UC+Oza), UC patients not treated with ozanimod (UC), or healthy controls (HC), and analyzed by flow cytometry. (E) Frequency of AnnexinV+ cells among naïve CD4+ T cells. Compilation of 10 experiments with n=8-11 per group. (F) Representative histograms and compilations of pJNK and pMKK7 relative geometric MFI values for naïve CD4 T cells. Relative values represent expression in one individual sample divided by the mean value of samples from the indicated comparison group on that day. In the first column, UC patients not treated with ozanimod were compared to healthy controls. In the second column, UC patients treated with ozanimod were compared to healthy controls. In the third column, UC patients treated with ozanimod were compared to UC patients not treated with ozanimod. Each UC patient with ozanimod is represented only once on the graph; if there was a choice between comparing that patient with a healthy control or a UC patient not taking ozanimod, the UC patient treated with ozanimod was compared to the UC patient not taking ozanimod. Compilation of 10 experiments with n=3-7 per group. (G) Representative histograms and compilation of relative geometric MFI values for BCL-2 and PUMA. Compilation of 10 experiments with n=3-7 per group. Statistical analysis was performed using Student’s t test for Figures 7B-7D and one-way ANOVA with Tukey’s multiple comparisons test for Figures 7E-7G. *p ≤ 0.05, **p ≤ 0.01, ***p ≤ 0.001, ****p ≤ 0.0001, N.S. non-significant.

We next asked whether increased T cell death was also present in patients treated with S1PR1 modulators. An exciting new application of these drugs is in ulcerative colitis (UC), with the S1PR1 modulator ozanimod (Zeposia^TM^) first approved by the U.S. Food and Drug Administration in 2021. The mechanism-of-action is assumed to be inhibition of pathogenic T cell migration from lymph nodes to the colon, although much remains to be learned. We analyzed peripheral blood mononuclear cells from UC patients being treated with ozanimod (9) (Table 1, Figure S6A). We found an increased percentage of AnnexinV+ cells among naïve CD4+ T cells as well as naïve CD8+ and central memory CD4+ T cells in patients treated with ozanimod compared to healthy controls or UC patients on a different therapy (Figures 7E and S6B). We also detected increased activation of the JNK pathway and an imbalance of BCL-2 family members in T cells from ozanimod-treated patients compared to controls (Figures 7F-7G, S6C-S6D). These data suggest that loss of S1PR1 function in human T cells, as in murine T cells, not only disrupts their trafficking but also may lead to apoptosis.

### Chronic FTY720 treatment reduces follicular T helper and germinal center B cell numbers

In response to the SARS-CoV-2 vaccine, patients on S1PR1 modulators have lower levels of anti-spike IgG antibody and lower seroconversion rates compared to patients treated with most other types of immune suppressants(41). Interestingly, the strength of the antibody response has been reported to correlate inversely with the duration of drug treatment before vaccination(11, 12), while the effects of these drugs on lymphocyte trafficking should be rapid and sustained. We hypothesized that one mechanism for the reduced response is loss of the repertoire of naïve T cells due to apoptosis(42).

To test this hypothesis, we studied the response of FTY720-treated mice to a T cell-dependent model antigen, sheep red blood cells (SRBC), injected subcutaneously. We enumerated follicular T helper cells (Tfh) and germinal center B cells (GCB) in the draining LN 8 days after immunization. We compared vehicle, ‘short’ FTY720 treatment (first dose 1 day prior to immunization), and ‘long’ FTY720 treatment (first dose 3 weeks prior to immunization) (Figure 8A). Interestingly, the long FTY720 treatment led to a significant reduction in the number of both Tfh and GCB cells compared to other groups (Figures 8B-8D and S6E).

**Figure 8:**
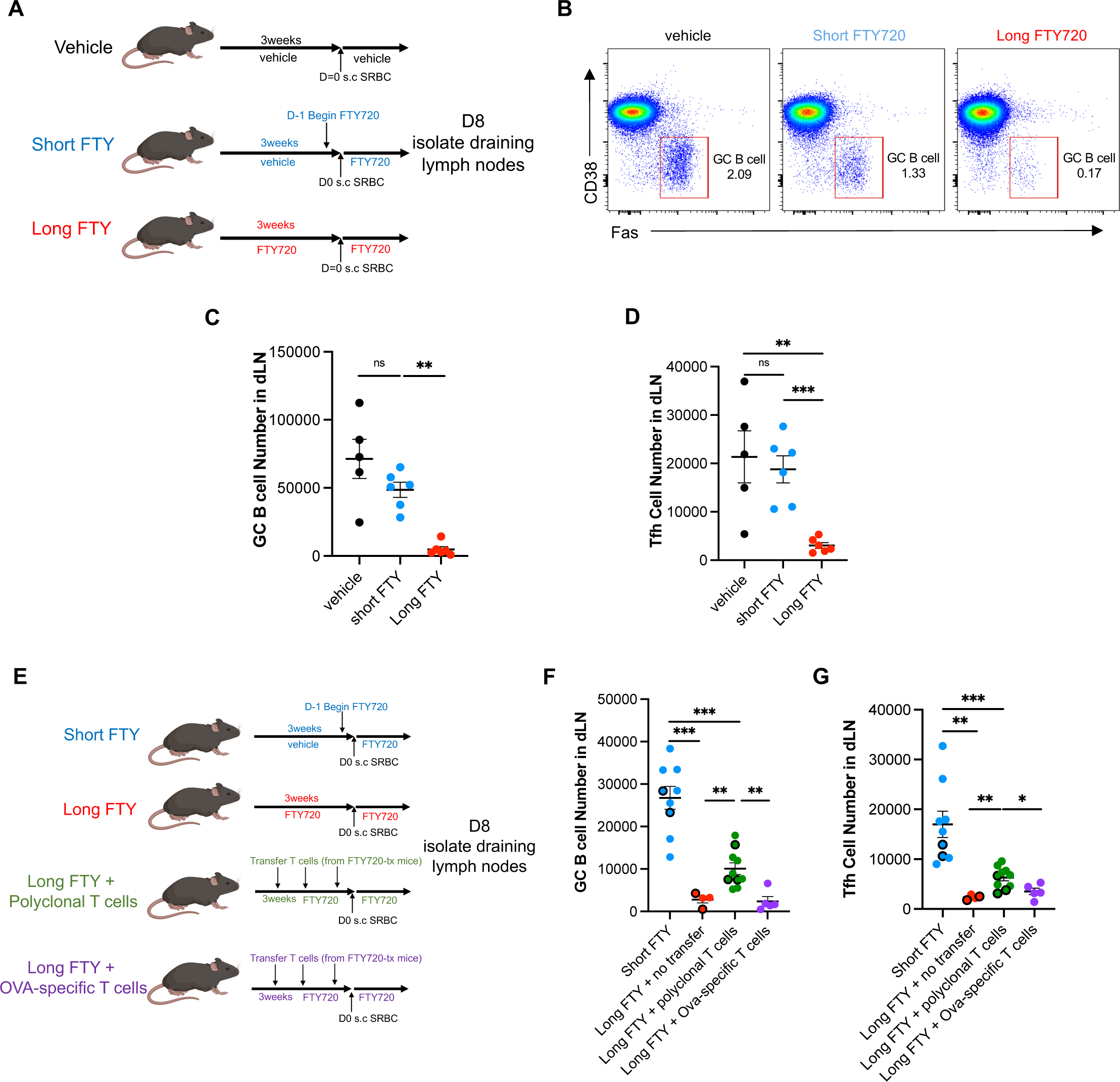
Prolonged treatment with FTY720 impairs germinal center responses in mice. A-D) WT mice were divided into 3 groups. The “long FTY” group received 1mg/kg FTY720 every other day for 3 weeks (starting on d −21). The “vehicle” group received vehicle for 3 weeks (starting on d −21). The “short FTY” was left untreated until d −1, when they received 1mg/kg FTY. On d0, all mice were subcutaneously injected in the left footpad with 10^8^ SRBCs. FTY720 (long and short FTY720 groups) or vehicle (vehicle group) treatment was continued until d8, when draining lymph nodes were analyzed. (A) Experiment design. (B) Representative flow cytometry plots of germinal center B cells (Fas+CD38-), gated on B220+ B cells. Numbers represent percent GC B cells of total B cells. (C) Compilation of number of GC B cells in draining lymph nodes. (D) Compilation of number of T follicular helper cells in draining lymph nodes. Compilation of 3 experiments with n=5-6 per group. E-G) Three groups of WT mice were treated with FTY720 for three weeks before immunization with SRBC (long FTY, treatment starting on d −21), while one group received FTY720 at d −1 before immunization with SRBC (short FTY). One of the long FTY groups (Long FTY + polyclonal T cells) received CD4+ and CD8+ polyclonal naive T cells from FTY720-treated WT mice (the donor mice initiated FTY720 at the same time as the recipient groups), either once (D −1: open dots) or three times (D −15, D −8, and D −1: closed dots). The second long FTY group received OVA-specific CD4+ and CD8+ T cells from FTY720-treated OTI and OTII transgenic mice (again, the donor mice initiated FTY720 at the same time as the recipient groups) three times (D −15, D −8, and D −1: closed dots); the ratio of OTI to OTII was calculated to match the ratio of CD8+ to CD4+ T cells in the polyclonal transfer. On D0, all mice were subcutaneously injected in the left footpad with 10^8^ SRBCs. FTY720 treatment was continued in all groups until D8, when draining lymph nodes were analyzed. (E) Experiment design. (F) Compilation of number of GC B cells in draining lymph nodes. (G) Compilation of number of T follicular helper cells in draining lymph nodes. Compilation of 4 experiments with n=5-10 per group. Statistical analysis was performed using Student’s t test. *p ≤ 0.05, **p ≤ 0.01, ***p ≤ 0.001, ****p ≤ 0.0001, N.S. non-significant.

FTY720 targets multiple S1P receptors on numerous cell types(43, 44), any of which might contribute to the declining germinal center response over time. To test whether the reduced germinal center response was due to loss of the naïve T cell repertoire (which in FTY720-treated mice, as in patients, was due both to cell death and a loss of thymic exit), we asked whether we could restore the response by increasing the number of naive T cells (Figure 8E). We transferred polyclonal naïve T cells into FTY720-treated mice. As a control, we transferred the same number of ovalbumin-specific T cells into a second group of FTY720-treated mice. If the decreased germinal center response were due to a loss of the naïve T cell repertoire, ovalbumin-specific T cells should not restore it. The transferred cells were isolated from donors treated with FTY720 using the same protocol as recipients, to avoid unanticipated artefacts. Transfer of polyclonal T cells, but not ovalbumin-specific T cells, increased Tfh and GCB cell numbers (Figures 8F-8G). The rescue was incomplete, but we were unable to replenish the T cell pool fully. The number of GCB cells increased with the total number of T cells in the case of polyclonal T cell transfer (Figure S6F). As expected, transfer of ovalbumin-specific T cells did not result in a large increase in the total number of T cells, likely due to competition for a limited niche (Figure S6F)(45).

## Discussion

Naïve T cell survival is crucial to maintain a diverse repertoire of T cells, which enables a robust immune response to a diverse range of pathogens. Here, we addressed the mechanism by which S1P and S1PR1 support naïve T cell survival, and whether this pathway is disrupted in patients treated with S1PR1 antagonists. We found that S1P signaling through S1PR1 limits JNK activation, which in turn maintains the appropriate balance of BCL2 family members within the T cell, and thereby prevents apoptosis. Interestingly, T cells without S1P/S1PR1 signaling have differential expression of genes regulating endocytosis and the actin cytoskeleton compared to controls, and phosphorylation of the same residues of the S1PR1 C-terminus that enable ligand-induced internalization of S1PR1 also promotes S1PR1-dependent survival. FTY720, a functional antagonist of S1PR1 that is used extensively to treat patients with autoimmune disease, induces naïve T cell death via the same pathway as genetic deletion of SPNS2 and S1PR1. The loss of naïve T cells can limit generation of B cell responses.

Our work suggests that S1PR1 modulators currently used in the clinic to treat autoinflammatory diseases may induce naïve T cell apoptosis. Early studies aimed at elucidating the mechanism for the lymphopenia induced by FTY720 found evidence of increased apoptosis of cells treated with the drug(46–48). Notably, one study found evidence of increased activation of JNK(49), and another study found that BCL-2 overexpression prevented FTY720-induced apoptosis *in vitro*(47, 48). Together with a long series of studies implicating S1P in cell survival, these experiments suggested that cell death could account for the lymphopenia and immunosuppression(19, 50, 51). In contrast, *in vivo* studies demonstrated that FTY720 induced lymphopenia by inhibiting egress of lymphocytes into lymph(44). Our experiments echo the early findings seen *in vitro* with FTY720 treatment. FTY720’s effects on cell death *in vivo* may have been discounted due to its more readily observable effects on cell trafficking.

One key future direction is to explore how these findings should alter clinical practice. Our work suggests that it will be important to consider effects of S1PR1 modulators on immune responses within the lymph node, as well as the more obvious effects on immune responses in non-lymphoid tissues. Reductions in naïve T cell numbers may limit the initiation of immune responses, and it will be fascinating in future work to assess how chronic JNK signaling may affect T cell function. In some settings, this may be an unacceptable side-effect. In other settings, this may be beneficial, limiting the recruitment of new T cells into an immune response. Going forward, effects of targeting S1PR1 on the immune system must be assessed more holistically.

On the more basic side, a fascinating question is how S1PR1 limits JNK activation. Our data suggest that S1PR1 internalization (or the amino acids that enable internalization) is required to restrain JNK. We are considering three candidate mechanisms. First, it is possible that S1PR1 signals through a unique partner within endosomes or other intracellular compartment, which in turn limits JNK phosphorylation. Intracellular functions of GPCR are increasingly appreciated(52); for example, after ligand-induced internalization, GPR35 traffics to the outer mitochondrial membrane, where it regulates protein-protein interactions(53). Second, it is possible that when S1PR1 cannot be internalized, the internalization machinery is freed to associate with other receptors, which may result in JNK activation. For example, β-arrestin, a key regulator of GPCR internalization, has a MAP kinase docking site that can enable JNK activation in some settings(54, 55). Third, it is possible that loss of S1PR1 internalization itself induces cell stress, which in turn results indirectly or directly in JNK activation. Receptor internalization requires rearrangement of the actin cytoskeleton, and stabilization of actin filaments or alterations in actin binding proteins can induce JNK activation(56). Furthermore, S1PR1 has recently been shown to promote a unique form of bleb-based movement by utilizing Ezrin-Radixin-Moesin proteins, which form linkages between the plasma membrane and the actin network(26). Ezrin phosphorylation is associated with activation of MKK7 and JNK in B cells(57), suggesting the possibility that ezrin phosphorylation-dephosphorylation events may regulate actin dynamics and thereby affect cell survival. Interestingly, mice deficient in the actin regulators *Dock8*(58) or *Coro1a*(59, 60) have perturbed T cell homeostasis due to both migration defects and increased cell death. Lymphocytes must be able to deform in order to squeeze through endothelial barriers as well as to move within organs, and perhaps the same receptors that guide their migration keep the cytoskeleton flexible.

While role of S1PR1 in T cell egress from lymphoid organs is increasingly well understood, to the point at which four drugs targeting S1PR1 have been FDA-approved for treatment of inflammatory disease and more are in clinical trials, the role of S1PR1 in T cell survival has just begun to be explored. It will be essential to understand this pathway to better direct use of this powerful class of drugs.

## Materials and Methods

### Study Design

The goal of this study was to understand why T cells die without S1P signaling. Hypotheses were tested in mice *in vivo*, and findings in mice were compared to the phenotype of human white blood cells *ex vivo*. For mice, sample sizes balanced statistical robustness and animal welfare, and negative results should not be over-interpreted. No animals were excluded from analysis unless they were clearly sick (hunched, low body weight). These criteria were pre-established, and are standard in the laboratory. No specific method of randomization was used to allocate mice into groups, although sex-matched littermates were used when possible. For human samples, we analyzed blood from all consented patients who were treated with ozanimod at NYU Langone’s Inflammatory Bowel Disease Center over a period of 16 months, and consented controls who visited the clinic the same day. The order of sample collection and data acquisition was designed to avoid experimental bias: collection and processing of samples from control and knockout, as well as treated and untreated animals/people, were alternated. Investigators were not blinded, because there were no qualitative measurements. No outliers were excluded, and no experiment was excluded unless a positive or negative control failed. These criteria were pre-established, and are standard in the laboratory. The number of samples and number of experiments are indicated in each figure legend.

### Mice

C57BL/6J (WT; CD45.2), B6.SJL-*Ptprc^a^Pepc^b^*/BoyJ (CD45.1), *Spns2*^f/f^ (61), *S1pr1*^f/f^ (62), *Bax*^-/-^ (30), *Lyve1*-Cre(16), *UBC*-CreERT2(15), MHCII^-/-^ (63), *Cd69*^-/-^ (64), *Sphk1*^f/f^(65), *Sphk2*^−/−^(66), OT-I(67), and OT-II(68) mice have been previously described. All mice were bred in New York University School of Medicine animal facilities. Mice were on a C57BL/6 or, in some cases, a C57BL/6×129 background. In all cases, mice were compared to littermate controls. Mice were 7–20 weeks old at the time of analysis. Male and female mice were used depending on availability, as sex did not seem to affect the results. Mice were housed in specific pathogen-free conditions. All cages were on a 12:12-h light:dark cycle (lights on, 07:00) in a temperature-controlled and humidity-controlled room. Room temperature was maintained at 72 ± 2 °F (22.2 ± 1.1 °C), and room humidity was maintained at 30–70%. All animal experiments were performed in accordance with protocols approved by the New York University Grossman School of Medicine Institutional Animal Care and Use Committee.

### Human blood samples

Peripheral blood was drawn from anonymous, adult IBD patients being treated with Zeposia (ozanimod), adult IBD patients being treated with a different therapy, or healthy control adults under protocol: Mucosal immune profiling in patients with inflammatory bowel disease IRB #: S12-01137.

### Mouse treatments

Tamoxifen (Sigma-Aldrich) was dissolved in corn oil (Sigma-Aldrich) by shaking at 37°C for 3 hours, either at 20mg/mL or 40mg/mL. 2mg tamoxifen was administered intraperitoneally on 5 consecutive days or 6mg of tamoxifen was administered by gavage on 2 consecutive days. Unless otherwise indicated, analysis was 3-4 weeks after the last tamoxifen treatment. Efficacy of tamoxifen was checked by measuring depletion of blood naïve T cells in the case of *Spns2*^f/f^ *UBC*-CreERT2, where we saw naïve T cells in blood decline ∼20-fold. In the case of *S1pr1*^f/f^ *UBC*-CreERT2, we checked the efficacy of tamoxifen by measuring surface S1PR1 or surface CD69 by flow cytometry. For *S1pr1*, our deletion efficiency ranged from 80-95%. We excluded experiments where deletion was less than 75%. We did not exclude non-deleted cells from analysis except when we sorted cells; this reduced the magnitude of differences we found, but was the most straightforward approach and avoided assumptions.

For bone marrow chimera generation, recipients were lethally irradiated with two 5 Gy doses of x-ray irradiation separated by 3 h, and received 2 × 10^6^ bone marrow cells by intravenous injection. Chimeras were analyzed at least 8 weeks after transplantation.

For IL-7Rα blockade, mice received 0.4mg anti-IL-7Rα (Bio X Cell) or isotype control (Rat IgG2a) intraperitoneally every 4 days for three total treatments.

For transfers of *Bax*^-/-^ or littermate control T cells, T cells were isolated by negative selection using biotinylated antibodies against CD11b, CD11c, CD19, CD25, NK1.1, TCRγδ, and Ter119 and Magnisort Negative selection beads (Thermo Fisher Scientific) according to manufacturer’s instructions, then labelled with CellTrace Violet or CellTrace Yellow (Thermo Fisher Scientific), before transfer. Dyes were swapped between genotypes between experiments. 1 × 10^6^ – 2 × 10^6^ T cells per group were transferred via retro-orbital injection into recipient mice.

For 4-deoxypyridoxine-HCl (DOP) treatment, mice received 10 g/l sucrose plus 30 mg/l DOP (Sigma-Aldrich) or 10 g/l sucrose alone in the drinking water for three weeks.

For SEW-2871 treatment, SEW-2871 (Cayman Chemical) was dissolved in ethanol at 9mg/mL as stock solution. Stock solution was diluted in vehicle of 50% Tween 20-PBS. Solution was sonicated in a water sonicator for 10 minutes and then injected intraperitoneally at 10mg/kg daily.

For H-89 treatment, 40mg/kg H-89 (Selleckchem) was given by gavage in vehicle of 30% PEG400, 1% Tween80, 64% PBS every day for a total of 12 days.

For dexamethasone treatment, dexamethasone (Sigma-Aldrich) was injected intraperitoneally in vehicle of PBS at 2mg/kg. For irradiation treatment, adult (8-12 weeks old) C57BL/6J mice were exposed to 1 Gy x-ray radiation.

For ABT-199 treatment, 60mg/kg ABT-199 (Selleckchem) was given by gavage in vehicle of 45% PEG400, 5% Tween80, and 42% PBS.

For JNK inhibitors, 15mg/kg/day SP600125 (Selleckchem) or 20mg/kg/day JNK-IN-8 (MedChemExpress) was given intraperitoneally in vehicle of 40% PEG400, 5% Tween80, and 50.5% PBS every day for a total of 12 days.

For FTY720 treatment, mice were injected intraperitoneally with 1mg/kg FTY720 (Cayman Chemical) in vehicle of 0.5% DMSO, 4.5% 2-hydroxypropyl-b-cyclodextrin and 95% PBS every other day for three weeks or as indicated.

For sheep red blood cell (SRBC) immunization, SRBCs (Colorado Serum Company) were washed three times in PBS. After the third wash, SRBCs were enumerated using Beckman Coulter Multisizer 3 and resuspended at 10^8^ SRBCs in 30μL. 10^8^ SRBCs were subcutaneously injected into the left footpad. Eight days after immunization, germinal center B cells were enumerated from the draining lymph nodes (popliteal and inguinal).

### Cell preparation

For adoptive transfer experiments, lymphocytes from spleen and lymph nodes were isolated by mechanical disruption and filtration through a 70-μm cell strainer. Lymph nodes were combined cervical, brachial, axillary, inguinal and mesenteric, unless otherwise indicated. Lymphocytes were enumerated with a cell counter (Beckman Coulter Multisizer 3) set to detect nuclei between 3.5 and 10 μm. T cells were purified by magnetic bead enrichment (Thermo Fisher Scientific) and then retro-orbitally injected into mice.

For analysis by flow cytometry, lymphocytes were isolated from spleen or LN by mechanical disruption and filtration through a 70-μm cell strainer in isolation buffer (PBS with 2% FBS, 1mM EDTA). LN were combined cervical, brachial, axillary, inguinal and mesenteric, unless otherwise indicated. Lymphocytes were enumerated with a cell counter (Beckman Coulter Multisizer 3) set to detect nuclei between 3.5 and 10 μm.

For CellTrace dye labeling, cells were resuspended at 20 x 10^6^/mL concentration in PBS and dyes were added for a final concentration of 5uM. Cells were incubated for 10 minutes at room temperature and the reaction was stopped by adding PBS-FBS to a final concentration of 20% FBS. Labeled cells were washed two times in PBS before injecting into mice.

To generate bone marrow chimeras, bone marrow cells were flushed from long bones of the leg. Red blood cells were lysed using ACK lysis buffer (Gibco). Cells were enumerated and resuspended at concentration of 2 x 10^6^ bone marrow cells/100uL PBS.

For human cell analysis by flow cytometry, peripheral blood mononuclear cells (PBMCs) were isolated using Lymphoprep (StemCell Technologies) and Sepmate-50 (IVD) tubes (StemCell Technologies), according to manufacturer’s instructions. After PBMC isolation, cells were enumerated and analyzed by flow cytometry.

### In vitro culture with ABT-199 and Fas ligand

Cell suspensions were prepared from LNs of mice as described above. Lymphocytes were treated either with varying concentrations of ABT-199 (Tocris) or Fas ligand + Enhancer (Enzo Life Sciences) for 4 hours in a 37°C 5% CO2 incubator. After 4 hours, cells were stained for surface markers and then stained for AnnexinV (BioLegend) and Propidium Iodide in AnnexinV binding buffer (BioLegend) for 15 minutes at room temperature.

### Flow cytometry

Staining for S1PR1 was done on ice in PBS supplemented with 0.05% sodium azide, 1 mM EDTA and 0.5% FBS. Cells (2E6 in 25 microliters) were stained for 90 min with anti-mouse S1PR1 (74.4 μg/ml; R&D Systems), washed twice in buffer, stained for 45 min with anti-rat IgG-biotin F(ab′)2 (9.5 μg/ml; Jackson Immunoresearch), washed twice in buffer and stained with streptavidin coupled with APC and the other surface membrane antibodies.

To stain for active-Caspase and AnnexinV, single cell suspensions from LNs (2E6 cells in 25 microliters) were stained for surface markers with 10uM CaspACE FITC-VAD-FMK for 1 hour in PBS supplemented with 1 mM EDTA and 0.5% FBS on ice. Cells were washed once in previous buffer and once in AnnexinV Binding Buffer (Biolegend). Then cells were stained with 1:50 dilution AnnexinV (BioLegend) Pacific Blue or APC in AnnexinV Binding Buffer for 15 minutes at room temperature.

For intracellular staining, cells were first stained with Fixable Live/Dead Blue (Thermo Fisher Scientific) according to manufacturer’s instructions. Then, cells were stained for surface markers on ice for 30 minutes. Cells were fixed using eBioscience FoxP3 Fixation/Permeabilization kit for either 1 hour or overnight at 4°C. After fixation, cells were stained for intracellular markers for 1 hour. If secondary antibodies were required, this step was performed for 30 minutes.

For phospho-staining, tissues were mechanically disrupted in PBS to create a single cell suspension and immediately fixed in equal volume of 4% PFA for 15 minutes. Cells were washed in isolation buffer (PBS with 2% FBS, 1mM EDTA) and then stained for surface markers using antibodies with fluorophores that were previously tested to work on fixed cells [CD4 BUV395 (clone RM4-5), CD8 PE (clone 5.3-6.7), CD62L BV421(clone MEL-14), CD44 FITC (clone IM7), and CD25 BV650]. After surface staining, cells were permeabilized using eBioscience FoxP3 permeabilization kit for 1 hour. Cells were stained for phospho-antigens for 1 hour with primary antibodies and then stained using secondary antibodies for 30 minutes.

Flow cytometry data were acquired on an LSR-II flow cytometer (Becton Dickinson) and analyzed using FlowJo version 10 software package.

### RNA isolation and sequencing

RNA was isolated from FACS-sorted naïve CD4+ T cells using TRIzol LS (Invitrogen) followed by DNase I (QIAGEN) treatment and cleanup with an Rneasy Plus Micro kit (QIAGEN), and quantified on a 2100 BioAnalyzer instrument (Agilent Technologies, Inc.). The RNASeq library prep and sequencing were performed at the NYU Langone Genome Technology Center. 5-20 ngs of total RNA was used to prepare libraries using Trio RNA-Seq library prep kit (Tecan Genomics, Inc.) following manufacturer’s instructions. The protocol contains the following steps: DNase treatment to remove any DNA in the sample, first strand and second stand cDNA synthesis from the input RNA, single primer isothermal amplification (SPIA) of the resultant cDNAs, enzymatic fragmentation and construction of unique barcoded libraries, PCR library amplification (4 cycles were used) and an AnyDeplete step to remove rRNA transcripts. The Agencourt AMPure XP bead (Beckman Coulter) purified libraries were quantified using by qPCR and the size distribution was checked using Agilent TapeStation 2200 system. The libraries were subjected to paired-end 50 bp sequencing on NovaSeq 6000 sequencer (Illumina).

For the RNA sequencing analysis, sequencing reads were mapped to the mouse reference genome (GRCm38.85/mm10) using the STAR aligner (v2.5.0c). Alignments were guided by a Gene Transfer Format file. The mean read insert sizes and their standard deviations were calculated using Picard tools (v.1.126) (http://broadinstitute.github.io/picard). The read count tables were generated using HTSeq (v0.6.0), normalized based on their library size factors using DEseq2, and differential expression analysis was performed. The read per million normalized BigWig files were generated using BEDTools (v2.17.0) and bedGraphToBigWig tool (v4).

### Recombinant DNA and retroviral transduction of T cells

For overexpression of S1PR1 mutants, cDNAs (synthesized by GenScript) were cloned into the retroviral plasmid MIGR1. MIGR1 was a gift from Warren Pear (Addgene plasmid #27490). The mutations in ST10A were alanine substitutions for S336, S351, S353, S355, S358, S359, T371, S374, S375, and S380. Plat-E cell lines, transfected with the constructs using Lipofectamine 2000 (Thermo Fisher Scientific), were used to produce retrovirus. Before retroviral transduction, naïve T cells from *S1pr1*Δ lymph nodes were isolated by negative selection using biotinylated antibodies against CD11b, CD11c, CD19, CD25, NK1.1, TCRγδ, and Ter119 and Magnisort Negative selection beads (Thermo Fisher Scientific) according to manufacturer’s instructions. The cells were activated using anti-CD3ε and anti-CD28 cross-linking in the presence of 10 ng/mL IL2. T cells were transduced with virus at 24 hours and 40 hours post-activation. For transduction, cells were centrifuged at 30°C 2500rpm for 90 mins in the presence of 4ug/mL polybrene (Sigma-Aldrich). 64hr post T cell activation, activated T cells were removed from anti-CD3ε/anti-CD28 and IL-2 stimulation and changed to media containing 5ng/mL IL-7 and 10ng/mL IL-15 to generate T cells with central memory properties. T cells in IL-7 and IL-15 media were split 1:2 every day in new media containing IL-7 and IL-15, and cultured for a total of 3 days. T cells were counted and transferred into C57BL/6J WT recipient mice. We did not sort cells before transfer. Instead, in analysis of recipient mice, we identified transferred cells using GFP as a marker (from the MIGR1 vector), and we gated on cells with similar GFP expression for comparisons.

### Transmission electron microscopy

FACS-sorted naïve CD4+ T cells from LNs and spleen were pelleted at 400 g for 4 mins by swinging bucket Eppendorf 5810 into BEEM conical capsule tips (EMS, #69913-05), and fixed in 2% glutaraldehyde and 2% paraformaldehyde in 0.1M sodium cacodylate buffer (CB, pH 7.2) overnight at 4°C. The sample was processed following rOTO method(69, 70). In brief, after washing with 0.1M CB three times for 5 min each, the cells were post-fixed with 2% osmium tetroxide and 1.5% potassium ferrocyanide in 0.1M CB for 1.5 hours on ice. After five washes of 3 min each in ddH_2_O at room temperature, cells were placed in a filtered solution of 1% thiocarbohydrazide (TCH, Electron Microscopy Sciences, EMS, PA) in ddH_2_O for 20 min at room temperature to allow additional staining. Cells were washed five times in ddH_2_O again on ice, and placed in 2% aqueous OsO_4_ for an additional 40 min on ice. Finally, cells were washed an additional five times in ddH_2_O on ice, and placed in 1% aqueous uranyl acetate at 4°C overnight. Cells were washed five times in ddH_2_O at room temperature, and en bloc lead staining was performed to enhance membrane contrast as follows. A lead aspartate solution was made by dissolving 0.066 g of lead nitrate in 10 ml of 0.003 M aspartic acid. The pH was adjusted to 5.5 with 1N KOH, and the solution was placed in a 60°C oven for 30 min. The lead aspartate solution was filtered, and cells were stained at 60°C for 30 min, then washed five times in ddH_2_O at room temperature and dehydrated in a graded series of ice-cold ethanol solutions (30, 50, 70, 85, 95, 100, 100%; 10 min each). Cells were then washed in ice cold acetone two times for 10 min each, followed by two acetone washes of 10 min each at room temperature. The cells were infiltrated gradually with EMbed812 epoxy resin (EMS) at room temperature by placing in 50% resin/acetone for 4 h, and 70% resin/acetone overnight. The cells were embedded in fresh 100% resin, and placed in a 60°C oven for 72 h to allow resin polymerization. The EMbed 812 resin recipe was 5 ml of EMbed 812, 4 ml of DDSA, 2 ml of NMA and 0.3 ml of BDMA. Ultrathin sections of 70 nm were cut, mounted on 200 mesh copper grids, and imaged with Talos120C transmission electron microscope (Thermo Fisher Scientific) with Gatan (4k x 4k) OneView Camera (Gatan, Inc., Pleasanton, CA).

### Mitochondrial morphological analysis

Mitochondrial numbers and cross-sectional surface area were obtained using ImageJ by manually tracing only clearly discernible outlines of mitochondria on TEM images. Approximately 25-30 whole cell images were taken per sample at the same magnification for each group within an experiment. Cross sectional area is reported in squared micrometers and number of mitochondria were enumerated per cell.

### Quantification and statistical analysis

Statistical analysis was performed using GraphPad Prism v9. Data in graphs show mean value ± SEM. The p values were calculated using unpaired two-tailed Student’s t test or one-way ANOVA with Tukey’s multiple comparisons test. *p < 0.05, **p < 0.01, ***p < 0.001, ****p < 0.0001.

## List of Supplementary Materials

Supplementary Figure 1: SPNS2-derived S1P and S1PR1 prevent apoptotic death of naïve T cells

Supplementary Figure 2: CD69 regulates increased pAKT and pERK in S1PR1-deficient naïve T cells

Supplementary Figure 3: Expression of BCL-2 family proteins in the absence of S1P signaling

Supplementary Figure 4: BCL-2 family proteins regulate S1PR1-dependent naïve T cell survival

Supplementary Figure 5: JNK signaling regulates S1P-dependent naïve T cell survival

Supplementary Figure 6: S1PR1 modulators impair T cell survival in humans and germinal center responses in mice

Supplementary Table 1: Characteristics of ulcerative colitis patients treated with ozanimod

Supplementary Table 2: Materials used in the study

## Acknowledgements

D.D. designed and performed experiments, and wrote the manuscript; V.M.H., Y.S.Z., and M.O. performed experiments; J.E.C. designed and performed experiments; J.A. provided patient blood samples; and S.R.S. designed experiments and edited the manuscript.

We thank Dan Littman, Jason Cyster, and Samantha Lux for thoughtful comments on the manuscript; Jason Cyster (UCSF) for *Grk2*^f/-^CD4-Cre and control bone marrow; Ramin Herati (NYU) for help in analyzing human PBMC; and Alice Liang (NYU) for expert assistance in electron microscopy. This work was supported by NIH R01AI085166 (SRS); NIH R01AI123308 (SRS); NIH R01DK093668 (KC); NIH R01CA237264 (JEC); NIH K23DK124570 (JEA); a Blood Cancer Discoveries Grant from the Leukemia and Lymphoma Society, the Mark Foundation for Cancer Research and The Paul G. Allen Frontiers Group (SRS); Senior Research Award from the Crohn’s & Colitis Foundation (KC); a Clinical Investigator Research Award from the Crohn’s and Colitis Foundation (JEA); a pilot grant from the Judith & Stewart Colton Center for Autoimmunity (JEA); a Vilcek MSTP Scholar Fellowship (DD); NIH 5T32AI100853 (DD); NIH T32AR069515 (DD); NIH T32HD007520 (MO); NIH F31AI154793 (MO); and NIH P30CA016087, partially supporting NYU Grossman School of Medicine’s core microscopy, flow cytometry, and genomics facilities.

## Conflicts of Interest

S.R.S. is an inventor on provisional patent 63/413,443 (Inhibitors of SPNS2 and uses thereof). K.C. has received research support from Pfizer, Takeda, Pacific Biosciences, Genentech, and Abbvie. K.C. has consulted for or received an honorarium from Puretech Health, Genentech, Abbvie, GentiBio, and Synedgen. K.C. is an inventor on U.S. patent 10,722,600 and provisional patent 62/935,035, and 63/157,225. J.E.A. has received research support from BioFire Diagnostics, and consultancy fees, honoraria, or advisory board fees from BioFire Diagnostics, Janssen, Pfizer, BMS, and Abbvie.

## Data Availability

Further information and requests for resources and reagents should be directed to Susan Schwab. RNA-Seq datasets have been deposited in the NIH Gene Expression Omnibus as GSE221482.

---

**Supplementary Figure 1:**
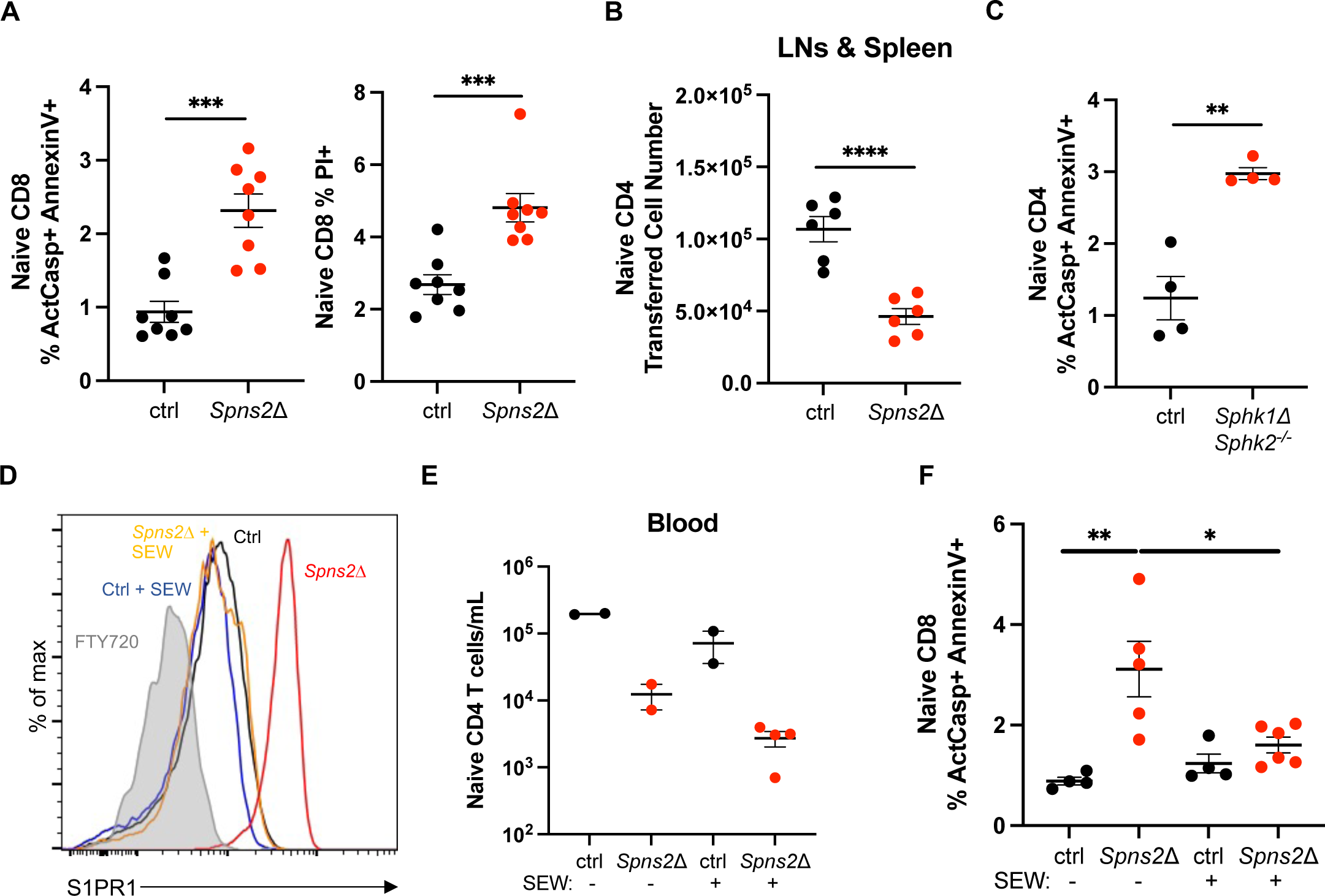
SPNS2-derived S1P and S1PR1 prevent apoptotic death of naïve T cells. A) *Spns2*^f/f^ *UBC*-CreERT2 mice and littermate controls were treated with tamoxifen, and 3-4 weeks later LN T cells were analyzed by flow cytometry. Frequency of ActCasp+ AnnexinV+ and PI+ cells among naïve CD8+ T cells. Compilation of 4 experiments with n=8 mice for both groups. B) *Spns2*^f/f^ *UBC*-CreERT2 mice and littermate controls were treated with tamoxifen. 3-4 weeks after tamoxifen treatment, the mice received 1×10^6^ CD45.1+ lymphocytes intravenously. Three weeks post-transfer, naïve CD4+ CD45.1+ T cells remaining in the recipient animals were enumerated (total reflects sum of cells in skin-draining LN, mesenteric LN, and spleen). Compilation of 3 experiments with n=6 mice per group. C) Frequency of ActCasp+ AnnexinV+ cells among naïve CD4+ T cells in LN of *Sphk1*^f/f^*Sphk2*^-/-^ *Lyve1*-Cre mice and littermate controls. Compilation of 2 experiments with n=4 mice for both groups. D-F) *Spns2*^f/f^ *Lyve1*-Cre mice and littermate controls were treated with 10mg/kg SEW-2871 or vehicle daily. After ten days of treatment, T cells were analyzed by flow cytometry. (D) Representative histogram of S1PR1 expression on naïve CD4+ T cells from LN of *Spns2*Δ mice and littermate controls, with and without SEW-2871 treatment. (E) Number of naïve CD4+ T cells in the blood of *Spns2*Δ mice and littermate controls, with and without SEW-2871 treatment. (F) Frequency of ActCasp+ AnnexinV+ cells among naïve CD8+ T cells in LN of *Spns2*Δ mice and littermate controls, with and without SEW-2871 treatment. Compilation of 3 experiments with n=3-4 mice per group. Statistical analysis was performed using Student’s t test. *p ≤ 0.05, **p ≤ 0.01, ***p ≤ 0.001, ****p ≤ 0.0001, N.S. non-significant.

**Supplementary Figure 2:**
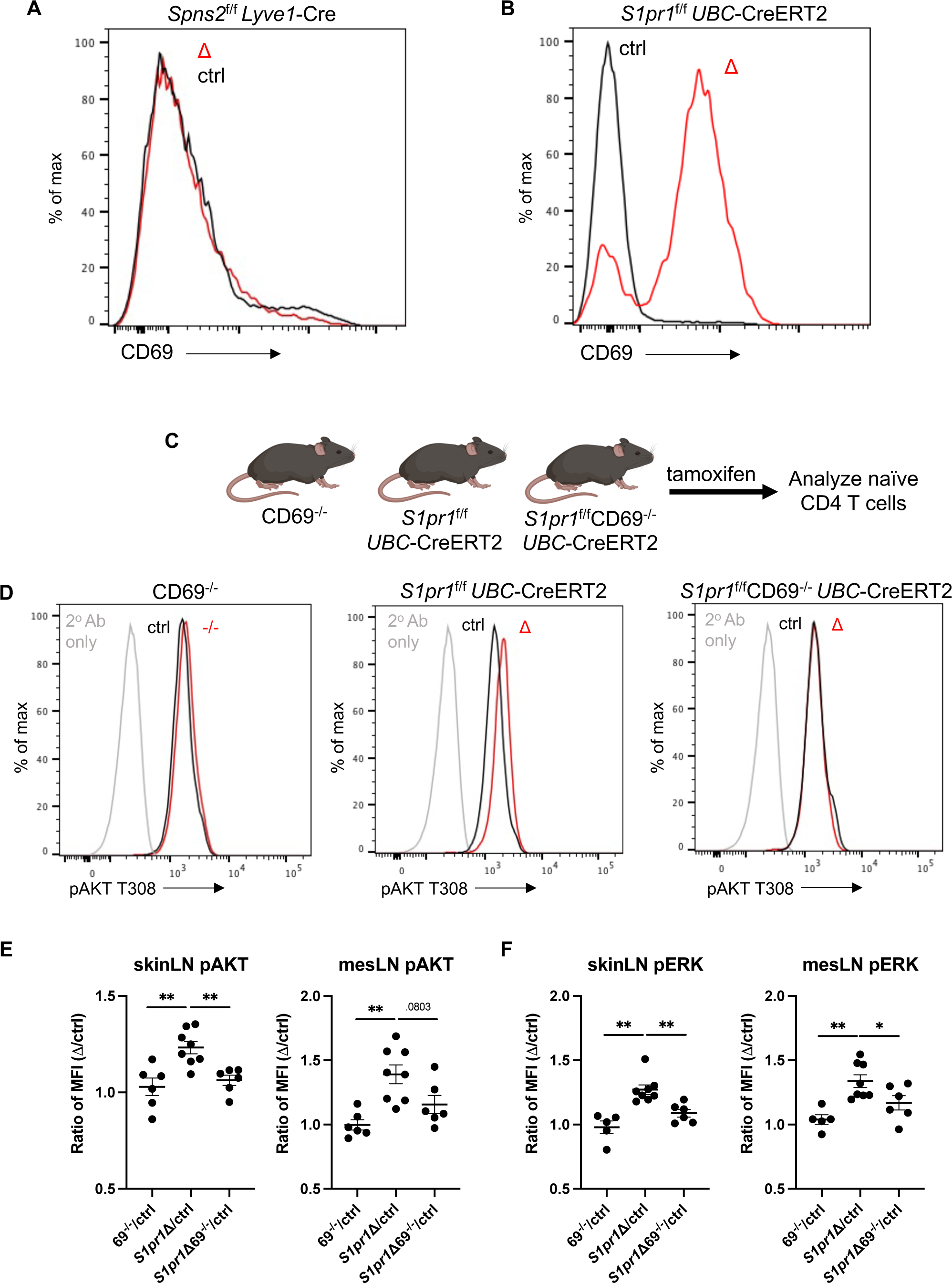
CD69 regulates increased pAKT and pERK in S1PR1-deficient naïve T cells. **Text:** One distinction between naïve T cells from *Spns2*Δ and *S1pr1*Δ mice is that naive *S1pr1*Δ T cells have high surface levels of the C-type lectin CD69 compared to controls, while naïve T cells from *Spns2*Δ mice do not have detectable surface CD69 (Figures S2A-S2B). CD69 and S1PR1 play antagonistic roles on the cell surface(71). Naïve T cells express high levels of S1PR1, which binds basal CD69 and maintains its low surface expression. A recently activated T cell robustly increases *Cd69* transcription, which shifts the balance; the increased CD69 binds S1PR1 and forces its internalization, keeping T cells at the site of activation. Naïve *S1pr1*Δ T cells have high levels of surface CD69 at baseline. Much remains to be learned about CD69’s function, but we hypothesized that surface CD69 might promote AKT and ERK signaling, as these pathways are normally induced early during T cell activation. To test this, we crossed *S1pr1*^f/f^*UBC*-CreERT2 with *Cd69*^-/-^ mice(64) and asked whether loss of CD69 reduced AKT and ERK phosphorylation in *S1pr1*Δ T cells (Figure S2C). *S1pr1*Δ naïve T cells had increased expression of pAKT and pERK compared to littermate controls and *Cd69*^-/-^ T cells, but *S1pr1*Δ; *Cd69*^-/-^ naïve T cells had similar expression of pAKT and pERK to controls and *Cd69*^-/-^ T cells (Figures S2D-S2F). This suggests that high expression of CD69 in *S1pr1*Δ naïve T cells partially explains the increased pAKT and pERK. **Legend:** A-B) Representative histogram of CD69 surface expression on naïve CD4+ T cells from LN of (A) *Spns2*Δ and (B) *S1pr1*Δ (red) or littermate control (black) mice. C-F) *S1pr1*^f/f^ *UBC*-CreERT2 mice, CD69^-/-^ mice, or *S1pr1*^f/f^ *UBC*-CreERT2; CD69^-/-^ mice and their respective littermate controls were treated with tamoxifen. 3-4 weeks later, T cells from either skin-draining LN or mesenteric LN were analyzed by flow cytometry for pAKT and pERK expression. (C) Experiment diagram. (D) Representative histograms of pAKT expression by naïve CD4 T cells from skin-draining LN of the indicated mice. (E-F) Compilation. Each point represents ratio of (E) pAKT or (F) pERK geometric MFI in naïve CD4+ T cells between a *S1pr1*Δ, CD69^-/-^, or *S1pr1*Δ CD69^-/-^ mouse and its respective littermate control. Compilation of 6 experiments with n=6-8 mice per group. Statistical analysis was performed using Student’s t test. *p ≤ 0.05, **p ≤ 0.01, ***p ≤ 0.001, ****p ≤ 0.0001, N.S. non-significant.

**Supplementary Figure 3:**
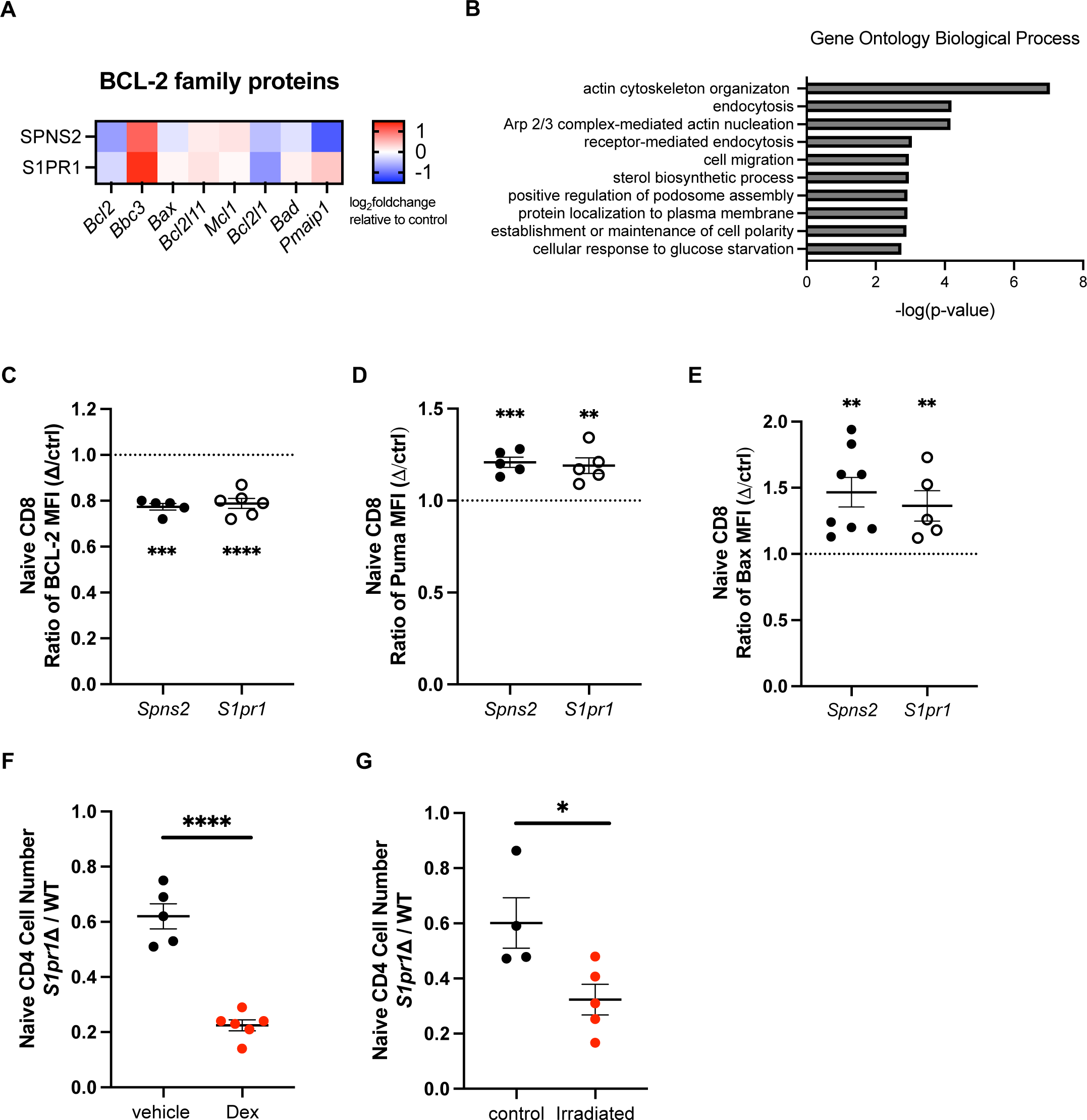
Expression of BCL-2 family proteins in the absence of S1P signaling. A-B) For SPNS2: CD45.2 *Spns2*^f/f^ *Lyve1*-Cre mice or CD45.2 littermate control mice were lethally irradiated and reconstituted with WT CD45.1/.2 bone marrow. After reconstitution, naïve CD4 T cells were isolated and RNA-Seq was performed. For S1PR1: a 50:50 mixture of CD45.2 UBC-GFP: CD45.2 *S1pr1*^f/f^ *UBC*-CreERT2 or CD45.2 *UBC*-GFP: CD45.2 *S1pr1*^f/f^ control bone marrow was transferred into lethally irradiated CD45.1/.2 WT mice. After reconstitution, mice were treated with tamoxifen. Four weeks after tamoxifen treatment, naïve T cells were isolated and RNA-Seq was performed. (A) Heatmap of BCL-2 family proteins in naïve CD4 T cells from *Spns2*Δ and *S1pr1*Δ mice. Heatmap represents log_2_fold change of genes from either *Spns2*Δ mice or *S1pr1*Δ mice relative to control cells. (B) Differentially expressed genes shared between both *Spns2*Δ and *S1pr1*Δ datasets were used to generate top 10 Gene Ontology (GO) biological process using DAVID Bioinformatics Resource. C-E) Expression of BCL-2 family members in naïve CD8+ T cells from LN of *Spns2*Δ or *S1pr1*Δ mice and littermate controls. (C) Compilation of BCL-2 expression. Each point represents the ratio of BCL-2 geometric MFI between a *Spns2*Δ or *S1pr1*Δ mouse and its littermate control. Compilation of 5-6 experiments with n=5-7 pairs per group. (D) As in (C) for PUMA. Compilation of 4-5 experiments with n=4-5 pairs per group. (E) As in (C) for BAX. Compilation of 6 experiments with n=5-8 pairs per group. F-G) *S1pr1*Δ and littermate control lymphocytes were co-transferred (1:1 by naïve CD4 T cell counts) intravenously into WT recipients. The transferred cells were labeled with CellTrace Violet or CellTrace Yellow, and dyes were swapped between experiments. 24 hours later, recipients were treated with either 2 mg/kg dexamethasone or 1 gy irradiation. 24 hours later, dye-labeled naïve CD4 T cells in LN were enumerated. Compilation of ratio of number of naïve CD4+ T cells from *S1pr1*Δ mice vs its littermate control recovered in recipients. (F) 2 mg/kg dexamethasone treatment. Compilation of 3 experiments with n=5-7 mice per group. (G) 1 gy irradiation. Compilation of 2 experiments with n=4-5 mice per group. Statistical analysis was performed using Student’s t test. *p ≤ 0.05, **p ≤ 0.01, ***p ≤ 0.001, ****p ≤ 0.0001, N.S. non-significant.

**Supplementary Figure 4:**
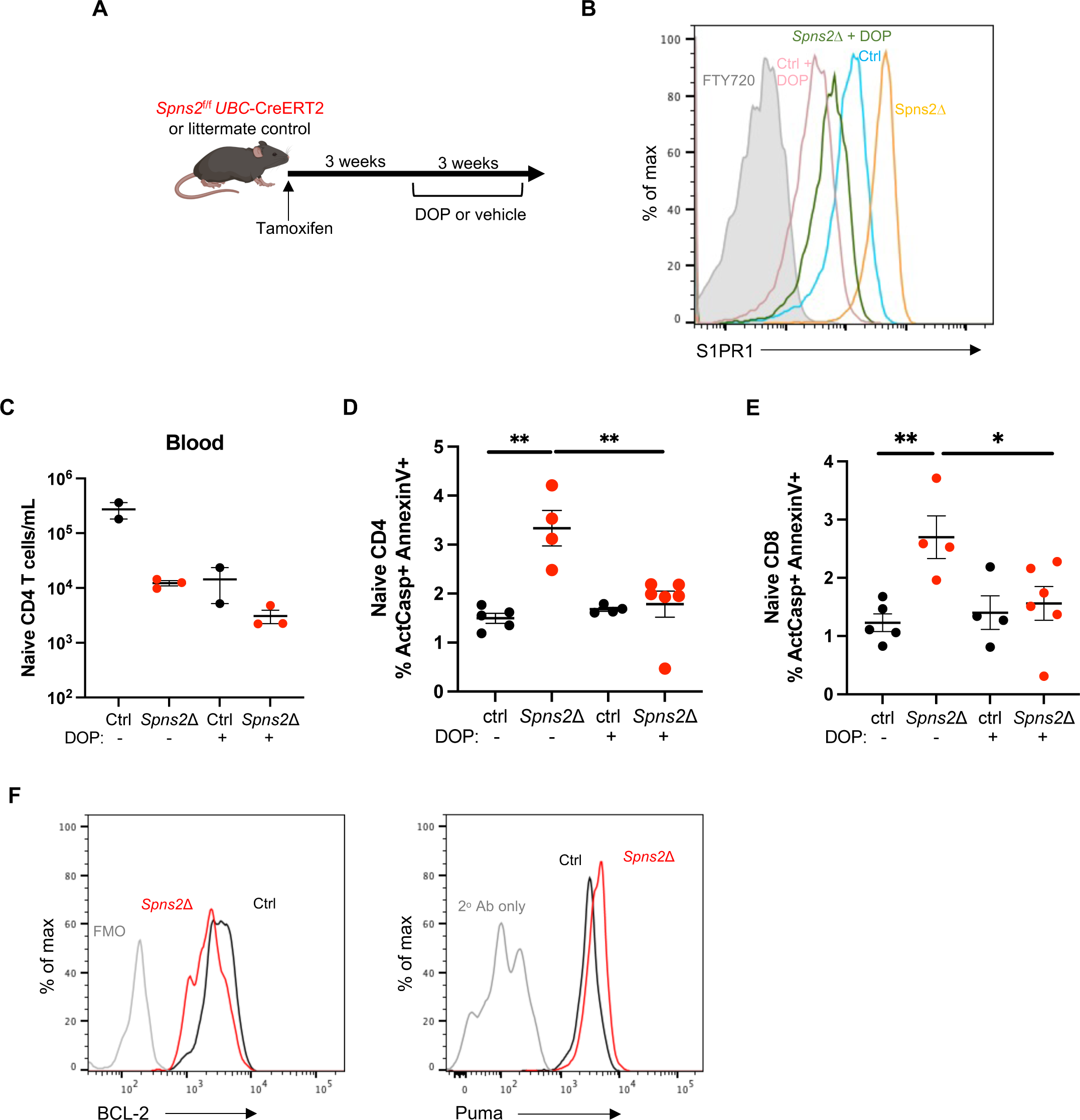
BCL-2 family proteins regulate S1PR1-dependent naïve T cell survival. A-E) *Spns2*^f/f^ *UBC*-CreERT2 mice and littermate controls were treated with tamoxifen. At least 3 weeks later, mice were treated with 30mg/l DOP and 10mg/l sucrose, or sucrose alone, in the drinking water. After 3 weeks of DOP treatment, LN T cells were analyzed by flow cytometry. (A) Experiment design. (B) Representative histogram of surface S1PR1 expression on naïve CD4+ T cells from LN of *Spns2*Δ mice and littermate controls, with and without DOP treatment. (C) Number of naïve CD4+ T cells in the blood of *Spns2*Δ mice and littermate controls, with and without DOP treatment. Compilation of 2 experiments with n=2-3 per group. (D) Frequency of ActCasp+ AnnexinV+ cells among naïve CD4+ T cells in LN of *Spns2*Δ mice or littermate controls, with or without DOP treatment. Compilation of 3 experiments with n=4-6 mice per group. (E) Frequency of ActCasp+ AnnexinV+ cells among naïve CD8+ T cells in LN of *Spns2*Δ mice or littermate controls, with and without DOP treatment. Compilation of 3 experiments with n=4-6 per group. F) Representative histograms of BCL-2 and PUMA expression in *Bax*^-/-^ naïve CD4+ T cells transferred to *Spns2*^f/f^ *Lyve1*-Cre mice or littermate controls. Statistical analysis was performed using Student’s t test. *p ≤ 0.05, **p ≤ 0.01, ***p ≤ 0.001, ****p ≤ 0.0001, N.S. non-significant.

**Supplementary Figure 5:**
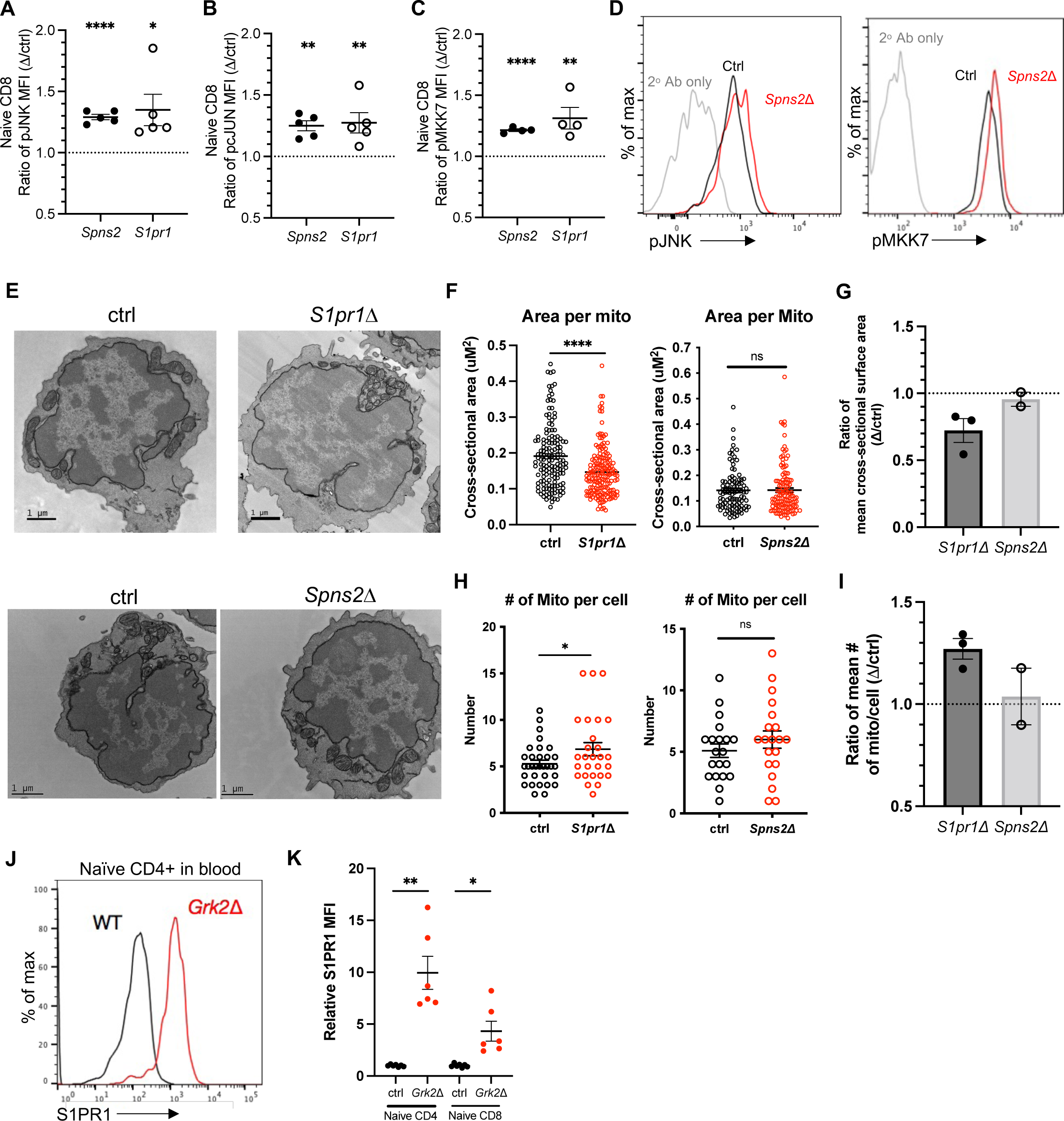
JNK signaling regulates S1P-dependent naïve T cell survival. A-C) Expression of phosphorylated JNK pathway proteins in naïve CD8+ T cells from LN of *Spns2*Δ or *S1pr1*Δ mice and littermate controls, analyzed by flow cytometry. (A) Compilation of relative MFI values for pJNK. Relative values represent expression in one individual mouse divided by the mean value of littermate control mice in that experiment. Compilation of 4-5 experiments with n=5 pairs per group. (B) As in (A) for pcJUN. Compilation of 4-5 experiments with n=4-5 pairs per group. (C) As in (A) for pMKK7. Compilation of 4 experiments with n=4 pairs per group. D) Representative histograms of pJNK and pMKK7 expression in *Bax*^-/-^ naïve CD4+ T cells transferred to *Spns2^f^*^/f^ Lyve1-Cre mice or littermate controls. E-I) Electron microscopy analysis of mitochondria in naïve CD4+ T cells sorted from LN. (E) Representative images of mitochondria within naïve CD4+ T cells from LN of a *S1pr1*Δ mouse and its littermate control or a *Spns2*Δ^Lyve1^ mouse and its littermate control. (F) Cross-sectional area of individual mitochondria, from one representative set of mice examining at least 20 cells per genotype. (G) Compilation of the ratio of the mean cross-sectional area of mitochondria in naïve CD4+ T cells of *S1pr1*Δ mice vs their littermate controls (3 pairs in 3 experiments) or of *Spns2*Δ mice vs their littermate controls (2 pairs in 2 experiments). (H) Number of mitochondria per cell, from one representative set of mice examining at least 20 cells per genotype. (I) Compilation of the ratio of the mean number of mitochondria per cell of *S1pr1*Δ mice vs their littermate controls (3 pairs in 3 experiments), or of *Spns2*Δ mice vs their littermate controls (2 pairs in 2 experiments). J-K) (J) Representative histogram and (K) compilation of surface S1PR1 expression on naïve T cells from blood of *Grk2*Δ and littermate control chimeras. Statistical analysis was performed using Student’s t test. *p ≤ 0.05, **p ≤ 0.01, ***p ≤ 0.001, ****p ≤ 0.0001, N.S. non-significant.

**Supplementary Figure 6:**
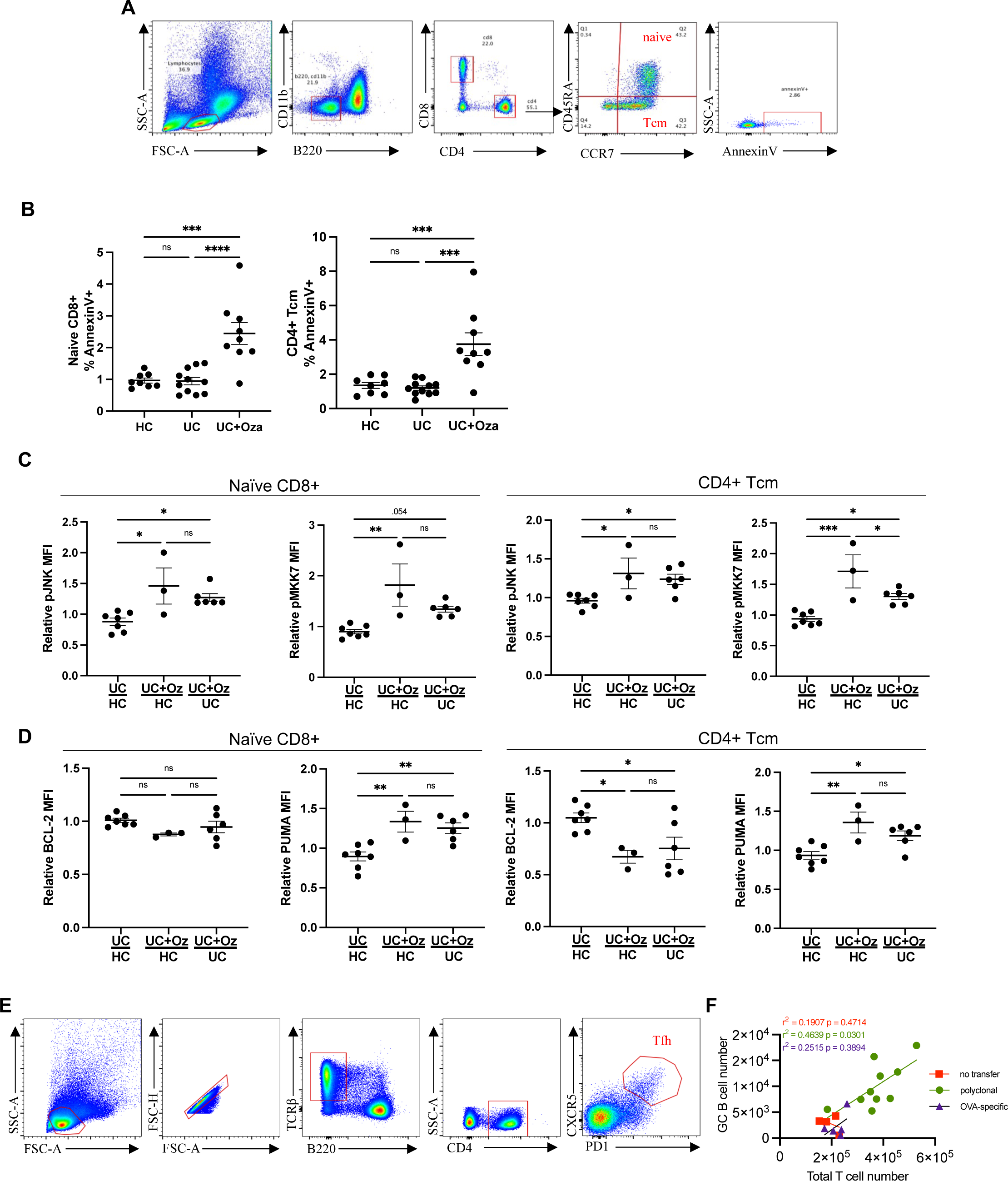
S1PR1 modulators impair T cell survival in humans and germinal center responses in mice. A-D) PBMCs were isolated from blood samples of UC patients treated with ozanimod (UC+Oza), UC patients not treated with ozanimod (UC), or healthy controls (HC), and analyzed by flow cytometry. (A) Gating strategy to analyze naïve CD4+, naïve CD8+, and CD4+ Tcm. (B) Frequency of AnnexinV+ cells among naïve CD8+ T cells or among CD4+ Tcm cells. Compilation of 10 experiments with n=8-11 per group. (C) Compilations of pJNK and pMKK7 relative geometric MFI values from naïve CD8+ T cells or from CD4+ Tcm cells, calculated as in Fig. 7F. Compilation of 10 experiments with n=3-7 per group. (D) Compilations of relative MFI values for BCL-2 and PUMA from naïve CD8+ T cells or from CD4+ Tcm cells. Compilation of 10 experiments with n=3-7 per group. (E) Gating strategy to identify Tfh cells. (F) Correlation between total T cell numbers and GC B cell numbers from ‘Long FTY + no transfer’ group, ‘Long FTY + polyclonal T cells’ group, and ‘Long FTY + OVA-specific T cells’ group. Statistical analysis was performed using one-way ANOVA with Tukey’s multiple comparisons test for Figures S6B-D. *p ≤ 0.05, **p ≤ 0.01, ***p ≤ 0.001, ****p ≤ 0.0001, N.S. non-significant.

**Supplementary Table 1:** Characteristics of ulcerative colitis patients and healthy controls.

|  | Healthy Controls (n=8) | UC+ No Ozanimod (n=11) | UC + Ozanimod (n=9) |
| --- | --- | --- | --- |
| Sex |  |  |  |
| Female | 5 (62.5%) | 2 (18.2%) | 4 (42.9%) |
| Male | 3 (37.5%) | 9 (81.8%) | 5 (57.1%) |
| Race |  |  |  |
| White | 6 (75%) | 6 (54.5%) | 5 (55.6%) |
| Black | 0 | 0 | 3 (33.3%) |
| Asian | 0 | 2 (18.2%) | 1 (11.1%) |
| Other/mixed | 2 (25%) | 3 (27.3%) | 0 |
| Hispanic ethnicity | 0 | 1 (9.1%) | 0 |
| Age at Encounter (years; median, IQR) | 32 (28) | 33 (17) | 29 (7.5) |
| Age at IBD diagnosis (years; median, IQR) | - | 26 (15) | 24 (7) |
| Duration of disease (years; median, IQR) | - | 6 (10) | 4 (4) |
| Maximum ulcerative colitis extent | - |  |  |
| Proctitis |  | 1 (9.1%) | 2 (22.2%) |
| Left-sided |  | 0 | 3 (33.3%) |
| Extensive colitis |  | 9 (81.8%) | 4 (42.9%) |
| Inflammatory biomarkers within 30 days | - |  |  |
| C-rp (median, IQR) |  | 0.7 (2.7) | 1.3 (4.6) |
| Fecal calpro (median, IQR) |  | 262 (250) | 839 (435) |
| Mayo endoscopic subscore within 30 days | - |  |  |
| 0 |  | 4 (36.3%) | 1 |
| 1 |  | 2 (18.2%) | 0 |
| 2 |  | 2 (18.2%) | 4 (42.9%) |
| 3 |  | 1 (9.1%) | 2 (22.2%) |
| Not assessed |  | 2 (18.2%) | 2 (22.2%) |
| IBD therapy exposure history | - |  |  |
| Mesalamines |  | 10 (90.9%) | 9 (100%) |
| Corticosteroids |  | 9 (81.8%) | 8 (88.9%) |
| Immunomodulators |  | 2 (18.2%) | 2 (22.2%) |
| Anti-TNF |  | 4 (36.3%) | 2 (22.2%) |
| Vedolizumab |  | 1 (9.1%) | 2 (22.2%) |
| JAK inhibitor |  | 0 | 2 (22.2%) |

**Supplementary Table 2:**
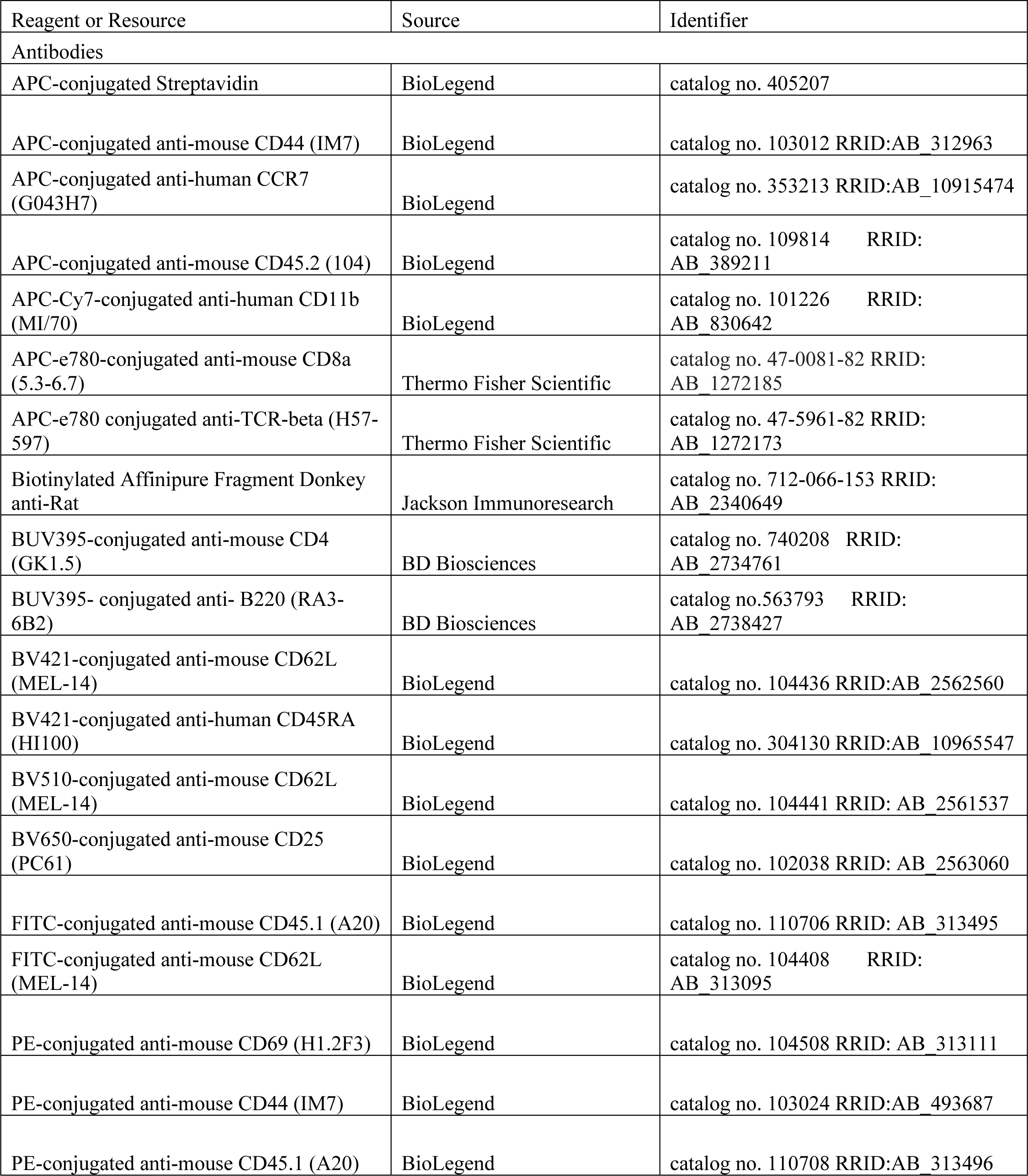
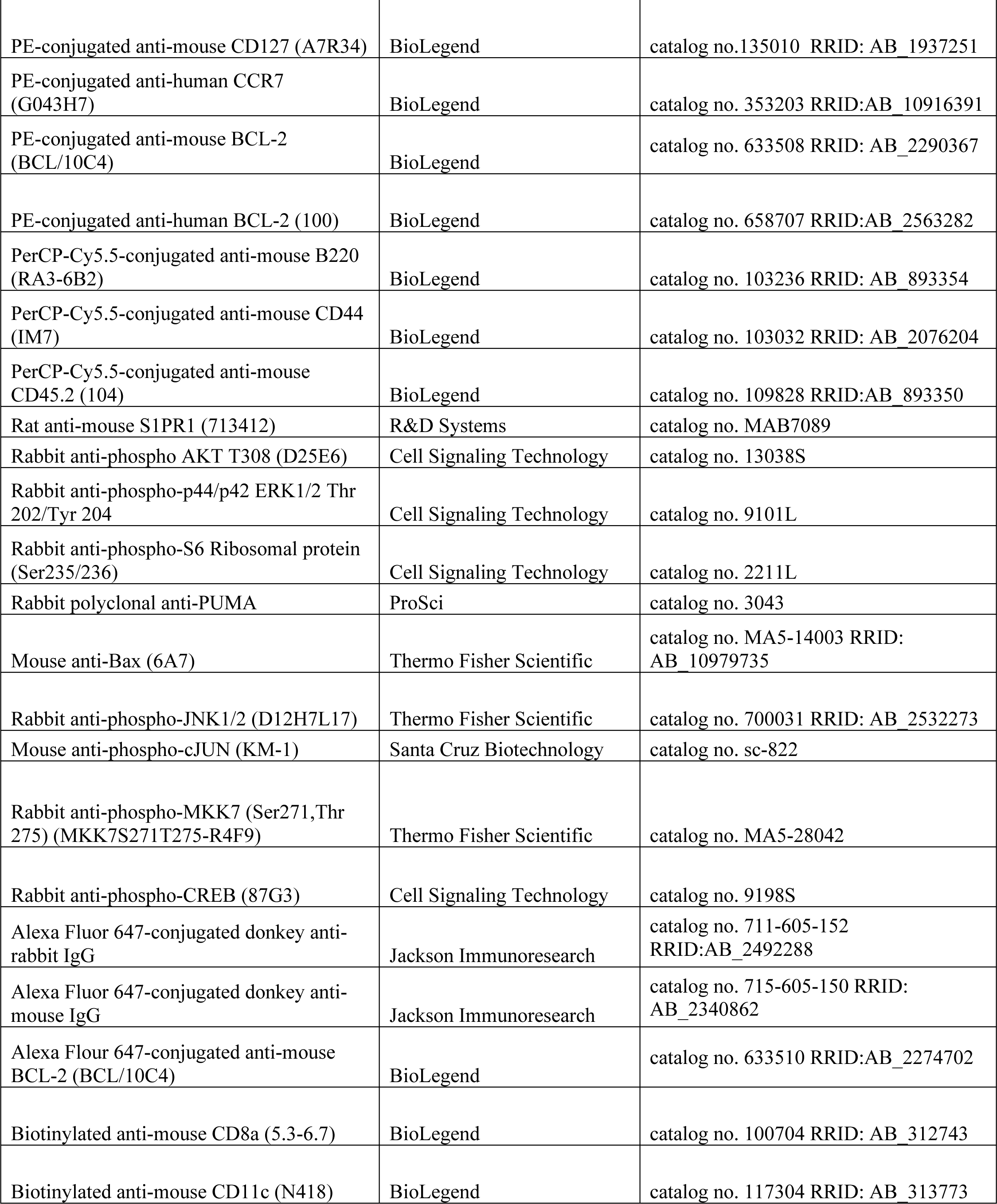
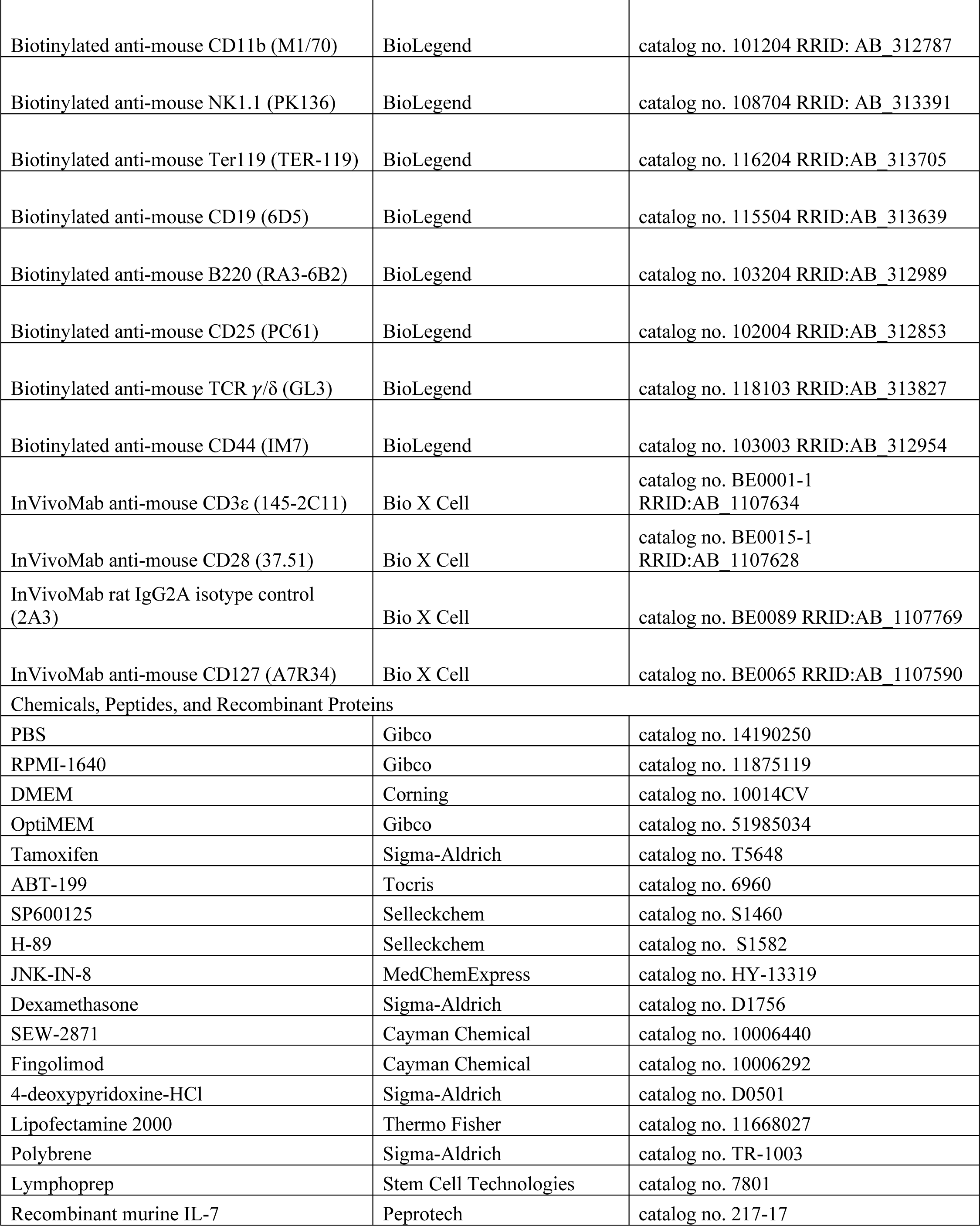
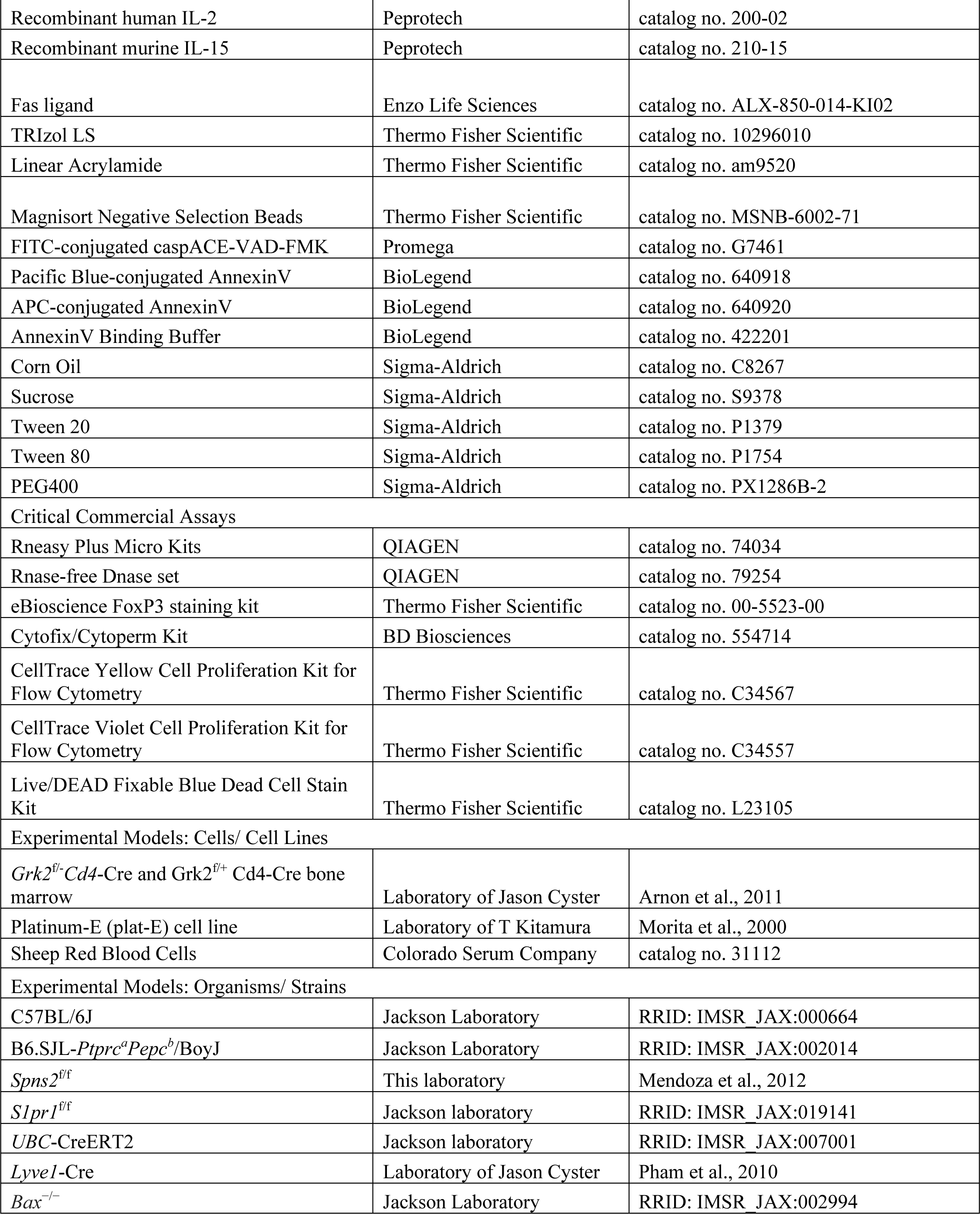
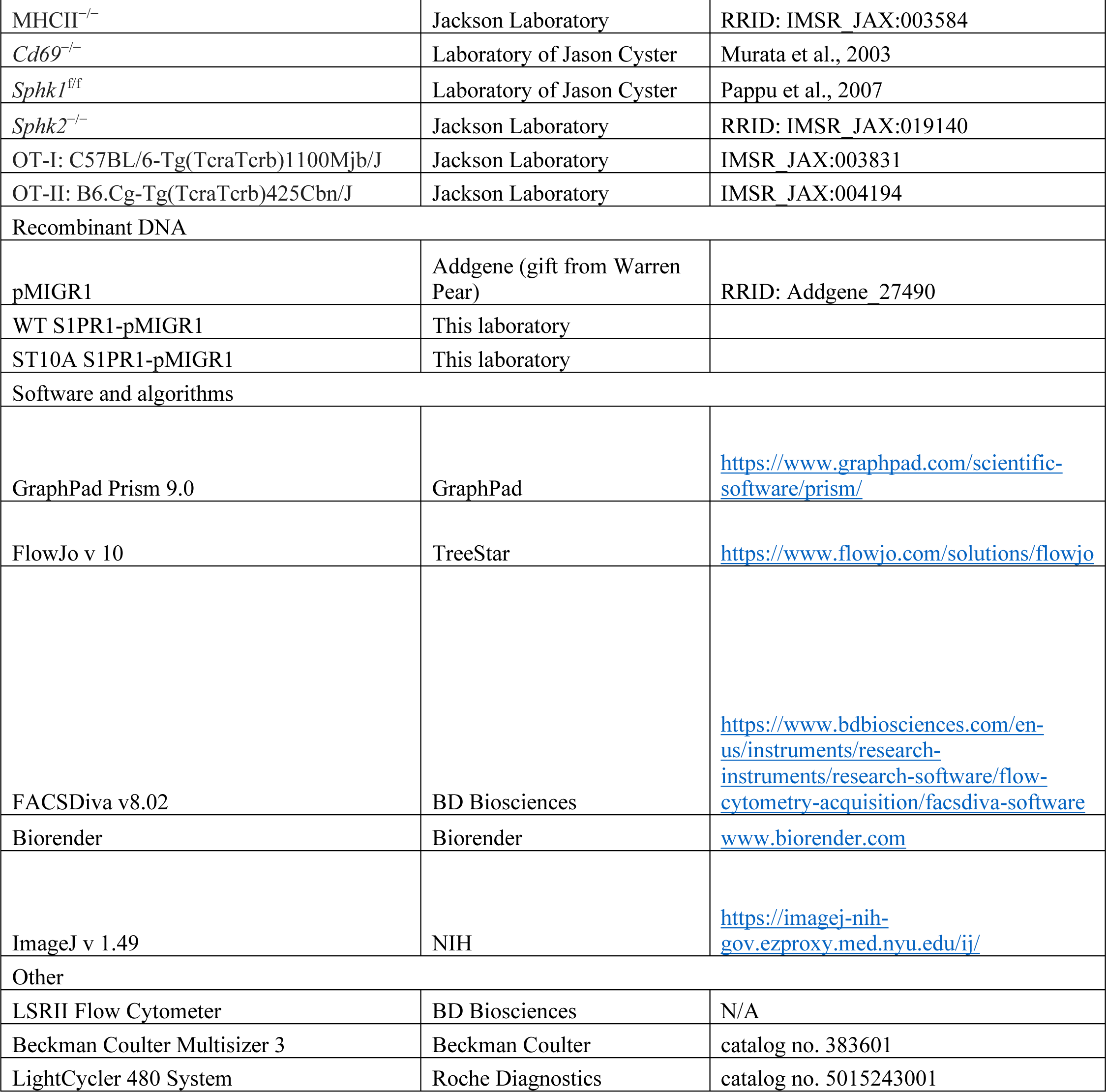
Materials used in the study.

